# 3D environment promotes persistent changes in lamin B1 distribution, the biomechanical signature of the nucleus, and adaptative survival and migratory functions

**DOI:** 10.1101/2023.04.10.536202

**Authors:** Raquel González-Novo, Héctor Zamora-Carreras, Ana de Lope-Planelles, Horacio López-Menéndez, Pedro Roda-Navarro, Francisco Monroy, Lin Wang, Christopher P. Toseland, Javier Redondo Muñoz

## Abstract

The interplay between cells and their surrounding microenvironment drives multiple cellular functions, including migration, proliferation, and cell fate transitions. The nucleus is a mechanosensitive organelle that adapts external mechanical and biochemical signals provided by the environment into nuclear changes with functional consequences for cell biology. However, the morphological and functional changes of the nucleus induced by 3D extracellular signals remain unclear. Here, we demonstrated that cells derived from 3D conditions conserve changes from cell confinement and show an aberrant nuclear morphology and localization of lamin B1, even in the absence of cellular confinement. We found that actin polymerization and protein kinase C (PKC) activity mediate the abnormal distribution of lamin B1 in 3D conditions-derived cells. These cells present altered chromatin compaction, gene transcription and cellular functions such as cell viability and migration. By combining biomechanical techniques and single-nucleus analysis, we have determined that the nucleus from 3D conditions-derived cells shows a different mechanical behavior and biophysical signature than the nucleus from control cells. Together, our work substantiates novel insights into how the extracellular environment alters the cell biology by promoting permanent changes in the chromatin, morphology, lamin B1 distribution, and the mechanical response of the nucleus.

## INTRODUCTION

The cell nucleus is organized into the nuclear envelope and the underlying lamina network, which encloses the chromatin and controls multiple DNA functions (1, 2). In the physiological context, cells have to respond to external stimuli, including biochemical signals, but also biomechanical properties of the extracellular microenvironment (3, 4). The mechanical response of the nucleus is dictated by the chromatin and the lamina network (5), which, in turn, is critical in multiple processes, including altered gene expression, cell fate transitions, cell migration, cell cycle progression, and tumorigenesis (6, 7).

Multiple biophysical techniques have been used to characterize the nuclear mechanics changes, such as cell stretching, substrate patterning, external compression, and 3D cell migration (8–10). Interestingly, studying the behavior of the nucleus in intact cells might lead to misinterpretation due to the cell membrane and the cytoskeleton; thereby, an integral study of intact cells and isolated nuclei might bypass this issue to better define the biophysical changes interpreted by the nucleus (11). This is particularly important from basic and translational research, as alterations in the deformability of the nucleus have been described in multiple human pathologies, including inflammation, aging, and cancer (12–14). It has been previously shown that three-dimensional (3D) *in vitro* confined environment compromises nuclear stability, promotes nuclear rupture and DNA damage, and perturbs different nuclear functions (15–18). However, it is not yet known how cell-environment interactions impact enduring alterations in the nucleus that drive genomic instability and cancer progression.

Here, we described how cell confinement endorses permanent changes that influence the mechanical and functional responses of the cell. We cultured cells embedded in a high-density 3D collagen-I matrix for several days, then collected and cultured them in suspension as TR (transformed) cells, which present permanent functional and phenotypical changes due to the memory of 3D confinement. In terms of morphological and phenotypical changes, we demonstrated that 3D confinement induced nuclear swelling and lamin B1 redistribution. Moreover, we identified that actin polymerization and protein kinase C (PKC) activity cause lamin B1 redistribution in altered cells. Mechanistically, we demonstrated that nuclear actin operates downstream of PKC and controls this change in lamin B1. Our transcriptional analysis reveals that transformed cells showed a different transcriptional activity and state compared to control cells, indicating potential differences between short-term cellular responses and the mechanical adaptation to 3D environments. Importantly, we revealed that 3D confinement affected the biology of transformed cells in functions such as migration, proliferation and survival. Finally, we combined atomic force microscopy (AFM), super resolution microscopy, confined compression, and optical tweezers, to demonstrate that transformed cells showed a homogeneous biomechanical signature inside the nucleus coupled to redistribution of chromatin. In summary, our results suggest that cell confinement can significantly impact nuclear changes, which could play a role in the regulation of other cell functions such as migration, transcription, and DNA repair.

## MATERIAL AND METHODS

### Cell culture

The human Jurkat (CVCL_0367) cell line was from ATCC, American Type Culture Collection. Jurkat cells were cultured in complete medium (RPMI 1640 medium with L-glutamine, 25 mM HEPES (Sigma), and 10% fetal bovine serum (Sigma)) and maintained in 5% CO_2_ and 37°C. TR cells were obtained upon long-term 3D confinement. For this, Jurkat cells were embedded in a solution of 3.3 mg/mL of bovine collagen type I (Stem Cell) in complete medium and neutralized with 7.5% NaHCO3 and 25 mM Hepes. After 1 hour at 37°C, additional complete medium was added at the top of the 3D collagen matrix. Jurkat cells were maintained in the 3D matrix for 10 days and the complete medium was changed every 3-4 days. Then, cells were collected from the collagen gel, cultured in suspension, and expanded as TR cells.

### Immunofluorescence

Cells were cultured in suspension, treated or not with specific inhibitors at 37°C. Cells were seeded onto 10 µg/mL poly-L-lysine-coated glass slides for 30 min. Then, cells were fixed with 4% formaldehyde for 10 min and permeabilized with 0.5% Triton x-100 (Tx-100) in PBS for 5 min. After 30 min blocking in 10% fetal bovine serum with 0.1% Tx-100 in PBS, samples were incubated with appropriate primary antibodies (1:100) for 1h at RT, followed by several PBS washes and 1h at RT incubation with secondary antibodies (1:200). Samples were stained with Hoechst 33342 1 µg/mL for 10 min at RT, washed with PBS and water, and mounted. Images were acquired on an inverted DMi8 microscope (Leica) using an ACS-APO 63x NA 1.30 glycerol immersion objective. Quantification and analysis of images were determined using ImageJ (National Institute Health). 3D reconstructions and videos were obtained by Leica software. For distribution analyses, lamin B1 signal from a cross-sectional plane of the cells was plotted and the mean of 15 cells was calculated and marked in red.

### Nuclear isolation

Nuclei were isolated by resuspending cells in buffer A (10 mM HEPES, 10 mM KCl, 1.5 mM, MgCl_2_, 0.34 M sucrose, 10% (v/v) glycerol, 1 mM DTT, 0.1% Tx100, and Roche protease inhibitor) for 5 min on ice. Following 3800 rpm centrifugation for 5 min, the pellet with the nuclei was resuspended in TKMC buffer (50 mM Tris pH 7.5, 25 mM KCl, 3 mM MgCl_2_, 3 mM CaCl_2_, and proteinase inhibitors) or PBS for conducting the indicated experiments.

### Atomic Force Microscopy (AFM)

AFM measurements were performed with a Bruker Bioscope Resolve mounted on an inverted microscope (Nikon Eclipse Ti2) connected to an ORCA-Flash4.0LT (Hamamatsu) camera. 8×10^5^ isolated nuclei from control and TR cells were seeded on 50 mm cell culture dishes (Willco GWST-5040) coated with 100 µg/mL poly-L-lysine following the protocol in (19, 20). Imaging was performed in filtered PBS at room temperature. Pre-calibrated Silicon Tip – Nitride cantilevers (PFQNM-LC-V2 Bruker) were used with a 70 nm tip radius. PeakForce Tapping/PeakForce Capture with a maximum indentation force of 0.5 nN was used. Indentation curves were fitted within Nanoscope Analysis (Bruker) using a cone-sphere model (21).

### Mechanical compression

Complete cells or isolated nuclei from cells were resuspended in PBS, dyed with Hoechst 33342 1 µg/mL and sedimented onto 10 µg/mL poly-L-lysine-coated plates and placed in the cell confiner device (4D cell). Following the manufacturer’s instructions, mechanical confinement was performed by pushing the nuclei with a glass slide with micropillars of 3 µm height. Images of at least 20 nuclei were taken with a 63x objective before and after the confinement by an inverted confocal DMi8 microscope (Leica) and analysis of the nuclear area was performed with ImageJ.

### Nuclear swelling stress

Isolated nuclei were resuspended in TKMC buffer and sedimented onto poly-L-lysine-coated plates. Nuclei were incubated or not with 5 mM EDTA or KCl for 10 min. Then, nuclei were fixed, permeabilized, and stained with Hoechst 33342 (1 µg/mL). Quantification and analysis of nuclear area were determined using ImageJ software.

### *In vivo* cell homing

NOD-SCID-Il2rg-/-(NSG) mice (*Mus musculus*), were bred and maintained at the Servicio del Animalario del Centro de Investigaciones Biológicas Margarita Salas (CIB-CSIC) with number 28079-21A. All mice were used following guidelines issued by the European and Spanish legislations for laboratory animal care. Control and TR cells were labelled with Cell Tracker Far Red (2 µM) and CFSE (5 µM), respectively, for 30 min; then, cells were mixed 1:1 and 5×10^6^ mixture cells were injected intravenously (IV) in the tail vein of 9 weeks-old NSG mice. Animals were euthanized 24 hours after injection, and the bone marrow from femurs, spleens, and livers were extracted, processed through mechanical disaggregation, and resuspended in erythrocytes lysis buffer (150 mM NH_4_Cl, 10 mM KHCO_3_, 0.1 mM EDTA). Samples were acquired in a FacsCanto II (Beckton Dickinson) cytometer, and the percentage of labelled cells infiltrated into the tissues was analyzed using FlowJo software.

## RESULTS

### Prolonged cell confinement in 3D conditions promotes permanent morphological changes in the nucleus

The size and morphology of the nucleus are critical for multiple homeostatic conditions, but also in human pathologies, such as cancer, where tumor cells show an abnormal morphology (22). First, we embedded leukemia cells (control) in 3D collagen gels and cultured them for 10 days. Then, cells were collected from the 3D environment and cultured in suspension without any further physical constriction as TR (transformed) cells (**Figure 1A**). Remarkably, the cellular transformation induced in TR cells remained permanent for several months and even after freezing-thaw cycles, suggesting a persistent memory of 3D confinement. We visualized the nuclear morphology of control and TR cells and found that TR cells showed bigger nuclei than their counterparts (**Figure 1B, C**). We also confirmed the aberrant morphology, characterized by diminished nuclear circularity in TR cells (**Figure 1D**). In addition to confocal analysis, the size and complexity of cells and nuclei can be determined by flow cytometry (23, 24), and we have confirmed that the distribution of isolated nuclei from TR cells presented a larger size and more complexity than those from control cells (**Figure 1E**). In order to further study the nuclear morphology, we visualized control and TR cells by electron microscopy (EM) and confirmed that TR cells showed an aberrant nuclear shape compared to control cells (**Suppl. Figure S1**). To determine whether TR cells might present different nucleolar disposition, we stained control and TR cells for nucleolin and found no differences in the number of nucleoli, although we found a significant increment in the intensity of the signal and the nucleolar area (**Figure 1F and Suppl. Figure S1**). Together, these results indicate that persistent confined environments alter the nuclear morphology and nucleolar state of TR cells.

**Figure 1.**
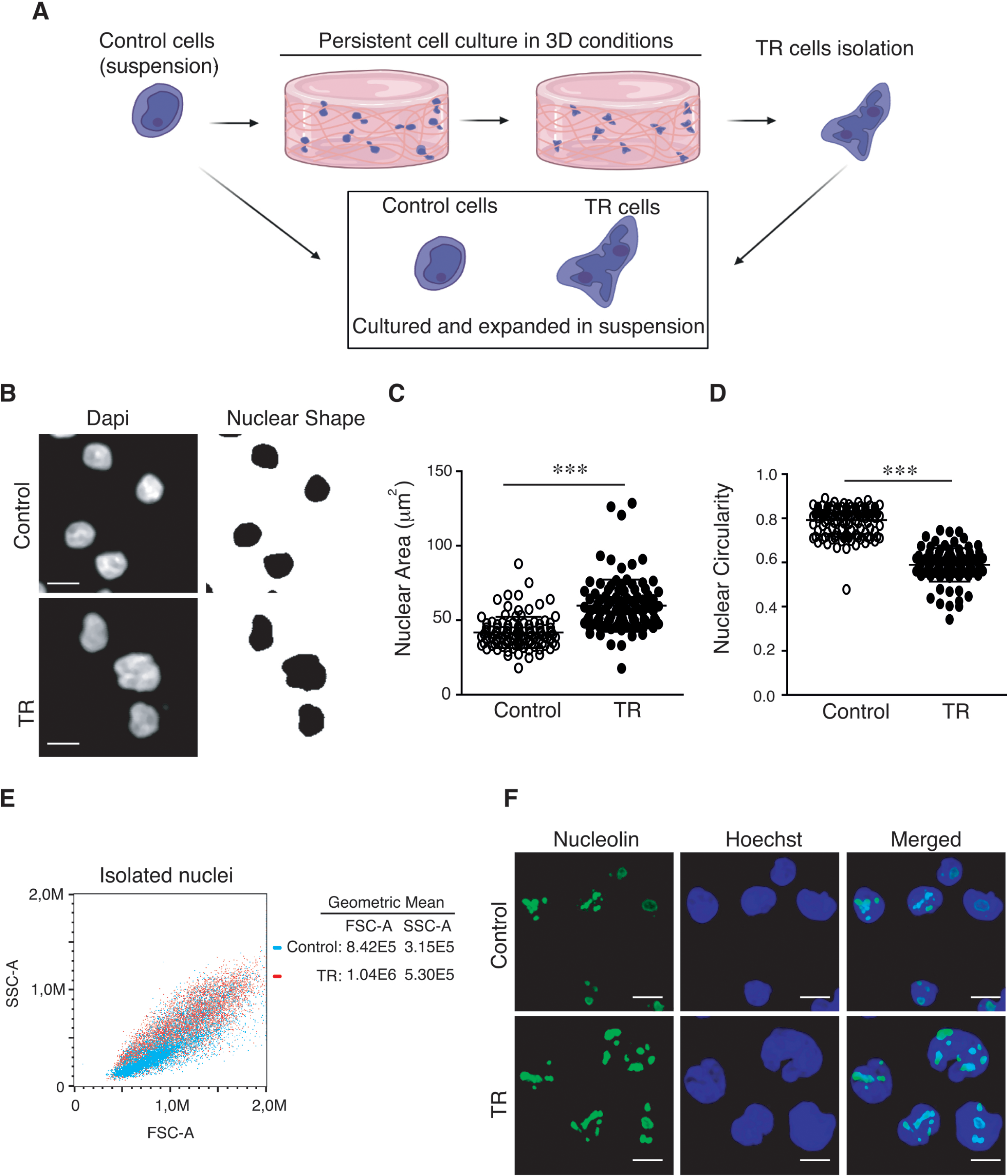
Persistent 3D cell confinement promotes permanent morphological changes in the nucleus. **(A)** Jurkat cells (control) were embedded in a 3D collagen matrix (3.3 mg/mL) in complete medium and maintained in culture for 10 days. Then, the collagen gel was disaggregated mechanically, and transformed (TR) cells were collected, kept in suspension without any further constriction, and expanded as TR cells. **(B)** Control and TR cells were sedimented on poly-L-lysine coated coverslips, fixed, and stained with Hoechst for their analysis by confocal microscopy. Right panels indicate the shape of the nuclei in black. Bar 10 µm. **(C)** Graph shows changes in the nuclear area of control and TR cells. Mean n=545 cells ± SD (3 independent replicates). **(D)** Graph shows changes in the nuclear circularity of control and TR cells. Mean n=545 cells ± SD (3 independent replicates). **(E)** Nuclei from control (blue) and TR (red) cells were isolated and their complexity (SSC-A) and size (FSC-A) determined by flow cytometry. Right panel indicates the geometric mean of both characteristics for both populations. **(F)** Control or TR cells were seeded on poly-L-lysine-coated glasses and stained with Hoechst (blue) and nucleolin (green) for their analysis by confocal microscopy. n=63-106 cells (3 independent replicates). Bar 10 µm. *** P<0.001.

### Persistent 3D cell confinement confers multilobular disposition of lamin B1 in the nucleus

Lamin B1 is the major component of the nuclear lamina of lymphocytes (25). Since 3D confinement alters the size and morphology of TR cells, we hypothesized that lamin B1 distribution may also be affected in these cells. First, we visualized the nucleus of control and TR cells and found that lamin B1 localized in the nuclear periphery of control cells, whilst TR cells presented an aberrant distribution of this protein (**Figure 2A-2C**). To distinguish whether aberrant lamin B1 distribution corresponded to intranuclear localization or multilobular reshaping of the nucleus, we characterized differences in the morphology and lamin B1 distribution of nuclei from control and TR cells by performing 3D reconstruction (**Movie S1, S2**) and by visualizing aberrant lamin B1 invaginations in middle cross-section plane of TR cells (**Figure 2D**). To confirm these results, we performed nuclear staining of the nuclear envelope marker, emerin, and found that most of the emerin signal localized at the nuclear lobes (**Movie S3, S4 Figure 2E**), suggesting that TR cells showed a multilobular shape of their nucleus. Lamin B type interacts and regulates nucleolar components (26–28), and we interrogated whether the aberrant distribution of lamin B1 in TR cells might regulate its interaction with nucleolar proteins. First, we performed immunoprecipitation experiments and found that TR cells presented similar levels of lamin B1 bound to the nucleolar protein NPM1 than control cells (**Suppl. Figure S2**). As expected, the disposition of nucleoli in TR cells was not connected to the abnormal distribution of lamin B1 (**Suppl. Figure S2**). These results indicate that TR cells showed an aberrant localization of lamin B1 in a multilobular shape.

**Figure 2.**
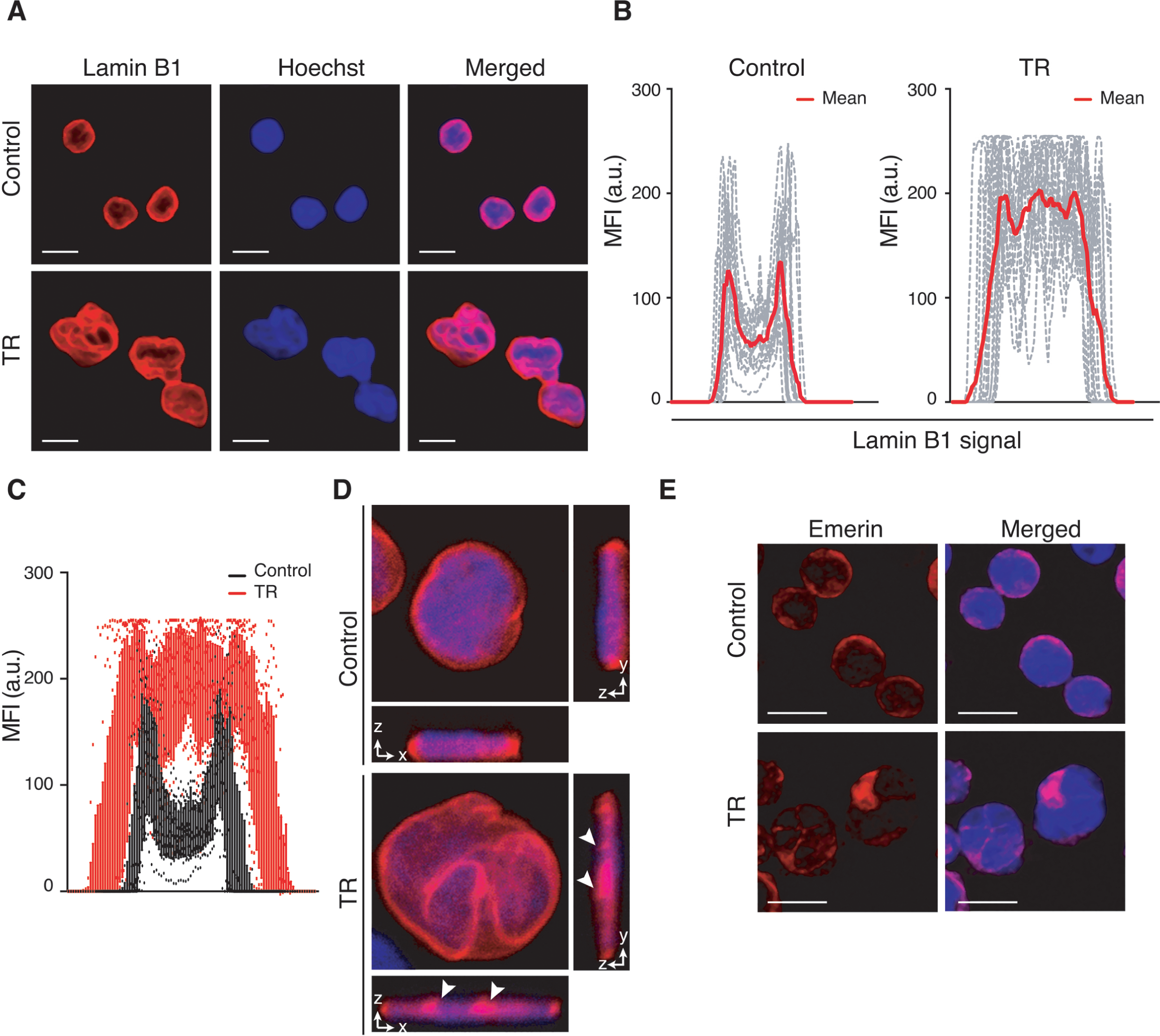
TR cells present multilobular nuclei characterized by an aberrant distribution of lamin B1. **(A)** Control or TR cells were seeded on poly-L-lysine-coated glasses and stained with Hoechst (blue) and anti-lamin B1 antibody (red) for their analysis by confocal microscopy. Bar 10 µm. **(B)** Line plots show the signal (MFI, mean fluorescence intensity) profile of lamin B1 for longitudinal sections from 15 representative cells from (A). Red line indicates the mean intensity of the profiles analyzed. **(C)** Graph shows the comparison between both populations (control and TR cells) for lamin B1 staining, resulting in a distinctive distribution of the signal intensity of their nuclei that resemble our previous observation. **(D)** Representative middle cross-section planes of control and TR cells stained as in (A). White arrows in orthogonal views indicate lamin B1 invaginations. **(E)** Control or TR cells were seeded on poly-L-lysine-coated glasses and stained with Hoechst (blue) and anti-emerin antibody (red) for their analysis by confocal microscopy. Bar 10 µm.

### PKC activity and actin polymerization modulate the aberrant distribution of lamin B1 induced by 3D confinement

It is well known that lamin B1 distribution is regulated by multiple kinases (29). We selectively inhibited conventional PKC activity with staurosporine (for PKCα) and enzastaurin (for PKCβ) and found that PKCβ inhibition suggested a partial recovery of the normal lamin B1 distribution in TR cells (**Suppl. Figure S3**). To rule out off-target effects of inhibitors and a complementary approach, we transfected TR cells with a specific pool of siRNAs against PKCβ (**Suppl. Figure S3**) and studied the distribution of lamin B1. As we anticipated, lamin B1 localized in the nuclear periphery in PKCβ-depleted cells (**Figure 3A, 3B**), confirming that PKC activity is required for this nuclear alteration induced by cell confinement. The cytoskeleton mediates mechanical signals between cells and their surrounding environment (30, 31). We addressed the potential implication of actin polymerization on the aberrant distribution of lamin B1 in TR cells. We incubated TR cells with chemical drugs to target actin polymerization and found that latrunculin B (an actin polymerization inhibitor) treatment promoted the localization of lamin B1 at the nuclear periphery whilst jasplakinolide, which stabilizes polymerized actin, did not significantly alter the distribution of lamin B1 (**Figure 3C, 3D**).

**Figure 3.**
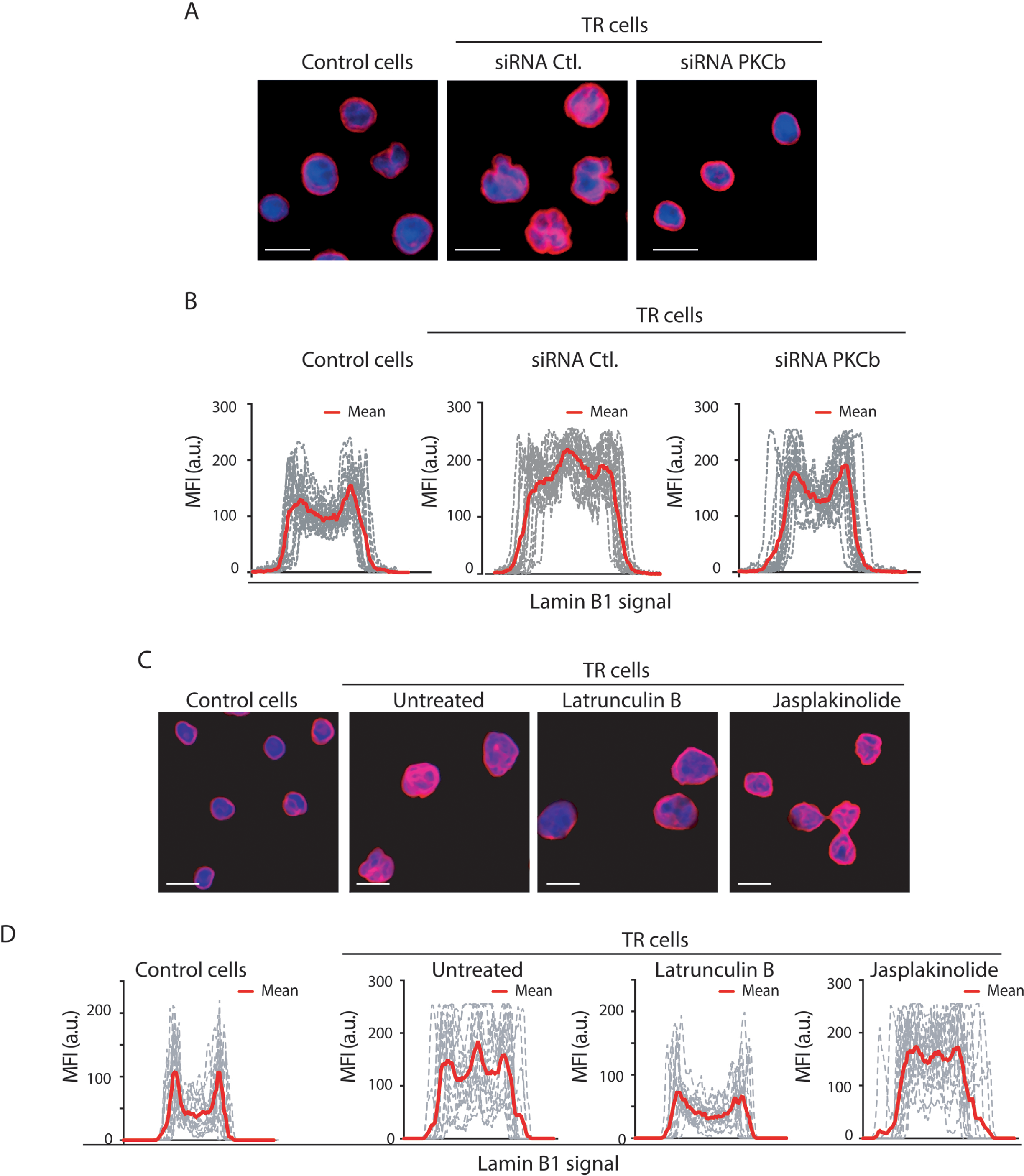
Persistent 3D cell confinement induces PKC-dependent redistribution of lamin B1 in TR cells. **(A)** TR cells were transfected with siRNA control or against PKCβ for 24 h. Then, cells were seeded on poly-L-lysine-coated glasses and stained with Hoechst (blue) and anti-lamin B1 antibody (red) for their analysis by confocal microscopy. Bar 10 µm. **(B)** Line plots show the signal profile of lamin B1 for longitudinal sections from 15 representative cells from 15 representative cells from (D). Red line indicates the mean intensity of the profiles analyzed. **(C)** TR cells were in the presence and absence of 2µg/mL latrunculin B (an inhibitor of actin polymerization) or 1µg/mL jasplakinolide (an actin polymerization stabilizer) for 1 h. Then, cells were seeded on poly-L-lysine-coated glasses and stained with Hoechst (blue) and anti-lamin B1 antibody (red) for their analysis by confocal microscopy. Bar 10 µm. **(D)** Line plots show the signal profile of lamin B1 from 15 representative cells from (A). Red line indicates the mean intensity of the profiles analyzed.

It has been reported that nuclear actin and its associated proteins play fundamental nuclear functions, such as chromatin changes, DNA repair, and transcriptional regulation (32). To further characterize the role of actin polymerization on the aberrant distribution of lamin, we transfected mutants of nuclear actin in TR cells and found that transfected cells with NLS-ActinR62D actin (an inactive mutant for actin polymerization) rescued in TR cells the phenotype observed in control cells. Nonetheless, the NLS-ActinWT and the NLS-ActinS14C (a mutant that recapitulates the effect of jasplakinolide) vectors did not significantly alter the distribution of lamin B1 (**Figure 4A, 4B**). As blocking actin polymerization and PKCβ signaling recovered the normal distribution of lamin B1, we interrogated which molecular mechanism was the primary signal controlling this nuclear change in TR cells. For this, we incubated TR cells with PMA (Phorbol 12-myristate 13-acetate) to stimulate the activation of PKC and observed that PMA treatment did not abrogate the effect of latrunculin on TR cells (**Suppl. Figure S4**). Furthermore, we treated TR cells depleted for PKCβ or preincubated with enzastaurin with jasplakinolide and found that actin stabilization overcame the recovery induced by targeting PKCβ (**Figure 4C, 4D**). When we inverted the order and pretreated TR cells with jasplakinolide before the addition of enzastaurin, we confirmed that actin stabilization also impaired the effect induced by PKC inhibition (**Suppl. Figure S4**). Taken together, these results suggest that nuclear actin polymerization regulates the distribution of lamin B1 in TR cells, and this effect might operate downstream the PKC signaling.

**Figure 4.**
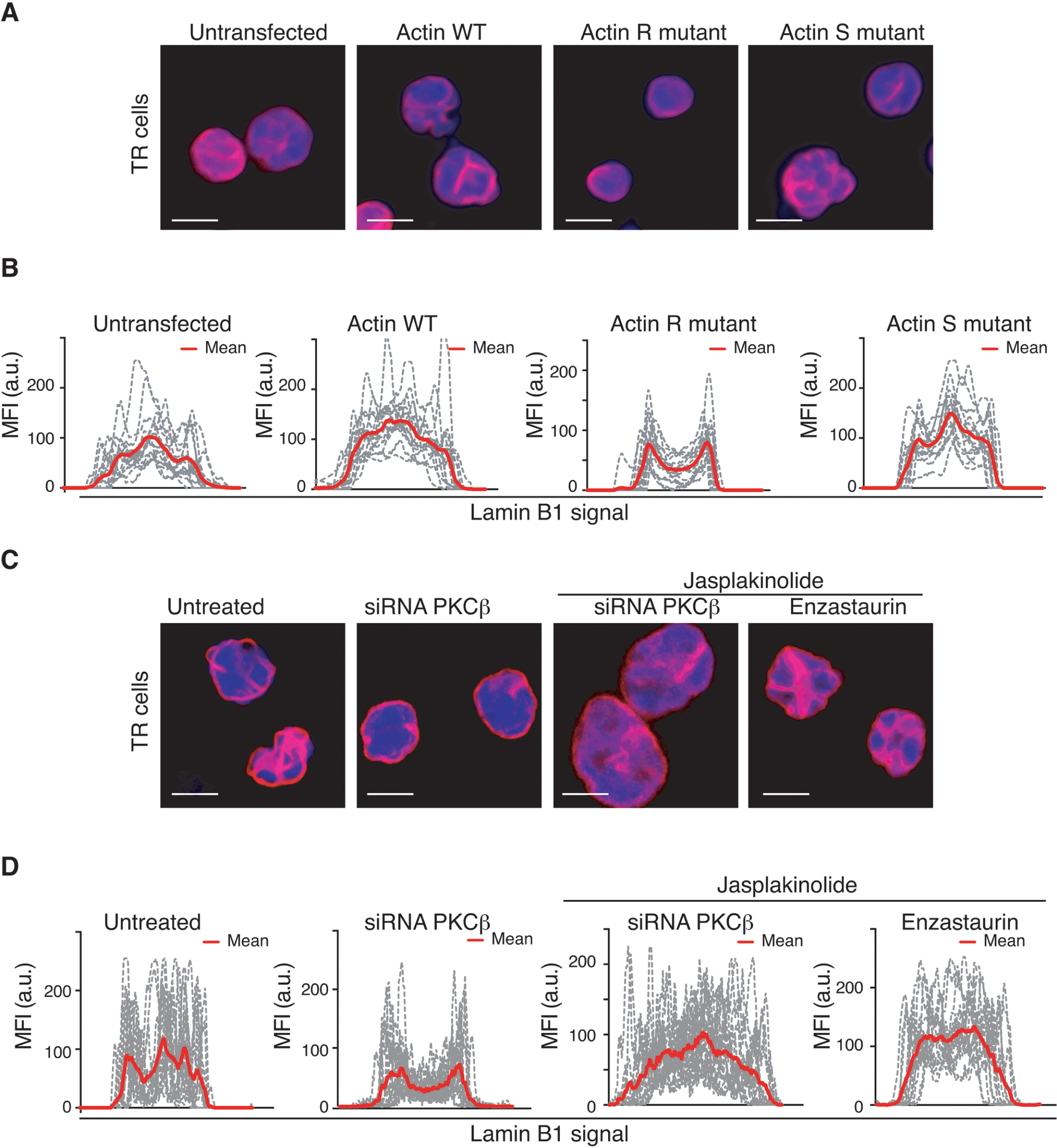
Nuclear actin polymerization modulates the aberrant distribution of lamin B1 in TR cells. **(A)** TR cells were transfected with different NLS-actin mutants for 48 h: NLS-YFP-WT (wild type actin), NLS-YFP-R62D (R actin to impair actin polymerization) and NLS-YFP-S14C (S actin to favour actin polymerization). Cells were seeded on poly-L-lysine-coated glasses and stained with Hoechst and anti-lamin B1 antibody (red) for their analysis by high content microscopy. Bar 10 µm. **(B)** Line plots show the signal profile of lamin B1 from 15 representative cells from (C). Red line indicates the mean intensity of the profiles analyzed. **(C)** TR cells were subjected to PKCβ silencing for 24 h or inhibition with 2 nM enzastaurin for 30 min, prior to treatment with 1µg/mL jasplakinolide for an additional hour. Then, cells were seeded on poly-L-lysine-coated glasses and stained with Hoechst and anti-lamin B1. Bar 10 µm. **(D)** Line plots show the signal profile of lamin B1 from 15 representative cells from (D). Red line indicates the mean intensity of the profiles analyzed.

### Aberrant cells after suffering 3D confinement alter their transcriptional program

It has been recently published that 3D collagen matrix density promotes cell fate transition in fibroblasts (33). As the actin cytoskeleton and an aberrant nuclear morphology are linked to the transcriptional activity of the cell (34), we interrogated whether these nuclear changes induced by 3D conditions were subsequent to gene expression alterations in TR cells. Our microarray expression analysis showed a significant differential expression of 1.4-fold change in 661 genes (143 up- and 518 down-regulated) in TR cells compared with control cells (**Figure 5A, 5B**). We performed bioinformatics analysis of the gene expression profiles in control and TR cells and found that the most significant changes correspond to genes encoding mitochondrial transport, cytokine signaling and extracellular matrix organization pathways (**Figure 5C, Suppl. Figure S5, and Table S1**). In order to validate our data, we selected two genes related to leukemia biology and determined their levels by RT-qPCR. Consistent with our transcriptional analysis, the levels of *MEN1* and *EHMT2* were diminished in TR cells (**Suppl. Figure S5**). Then, we assessed how 3D matrix environment might promote transcriptional changes at short-term in cells. For this, we cultured cells in suspension or embedded in a 3D collagen matrix for 3h, which has been reported to induce H3K4 methylation in leukocytes (16). Then, we analyzed their transcriptional activity observing that cells cultured at short times in 3D conditions also altered their transcriptional profile (**Figure 5D, 5E, Suppl. Table S2**). In spite of the experimental differences, we compared both 3D conditions and found 38 common genes altered in cells cultured for 3h in 3D conditions and in TR cells (**Suppl. Figure S5, Suppl. Table S3**); indicating that prolonged confinement promotes different outputs in the cell biology than short-term conditions. As we observed an increased intensity in nucleolin in TR cells, we interrogated whether these might show different transcriptional activity compared to control cells. To determine this, we measured the levels of RNA polymerase II and the incorporation of an alkyne-modified nucleoside, 5-ethynyl uridine (EU) in control or TR cells. Notably, TR cells showed increased levels of RNA polymerase II and EU than control cells, suggesting an increased transcription activity (**Figure 5F, 5G**). We confirmed in both cell populations the positive correlation between the RNA polymerase levels and the transcription activity, although control cells showed lower levels of these signals compared to the broad scatter in TR cells (**Figure 5H**). We further validated this increment of EU incorporation in TR cells by flow cytometry (**Suppl. Figure S5**). Together, these results demonstrate that persistent confinement in 3D environments promotes enduring changes in the transcriptional signature of cells.

**Figure 5.**
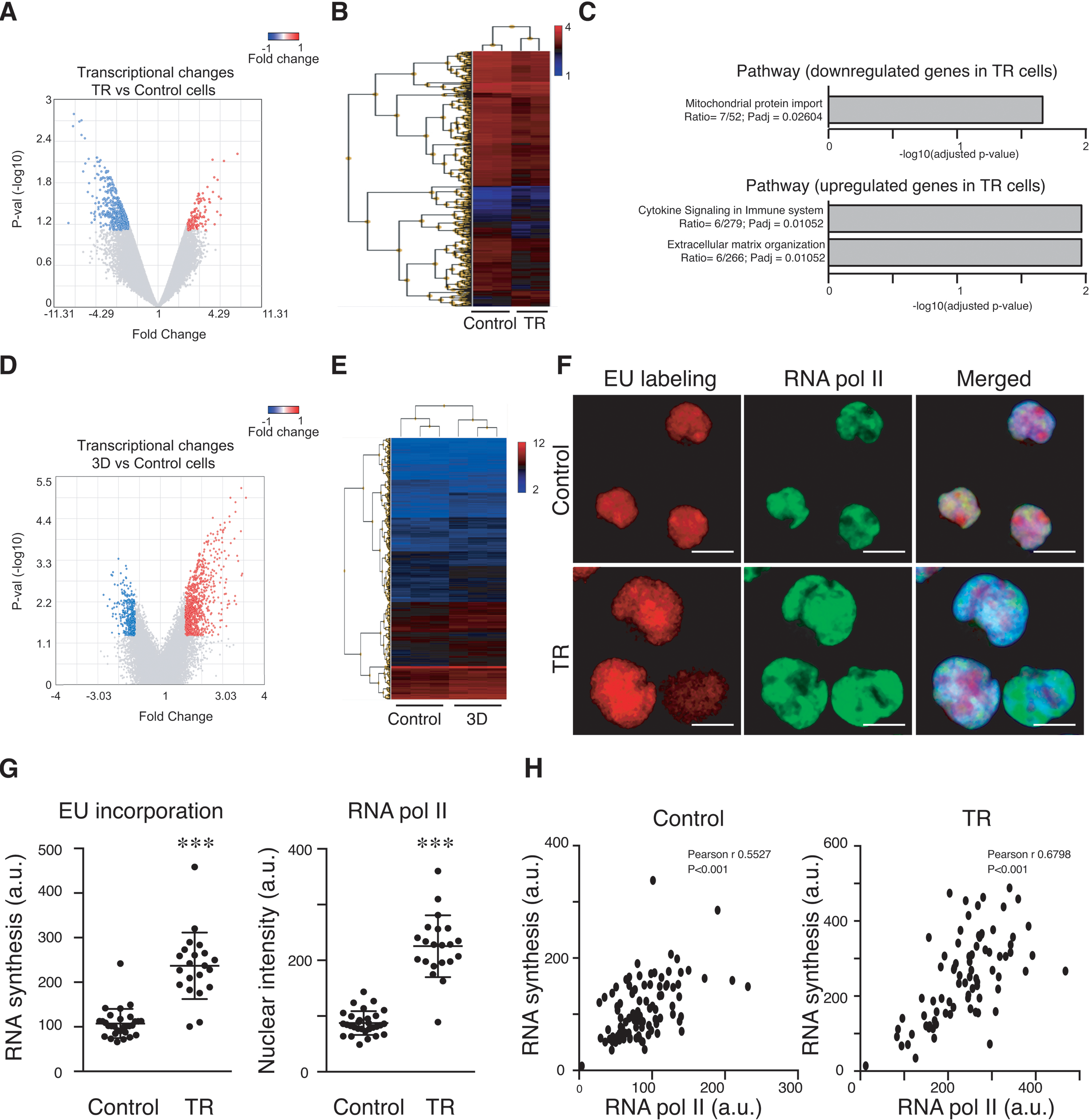
TR cells from 3D confinement alter their transcriptional program. **(A)** Control or TR cells were lysed, mRNA was isolated, and transcriptional changes were analyzed by mRNA expression microarray. Volcano plot shows significant downregulated (blue) and upregulated (red) transcripts in TR cells compared to control conditions. |Fold Change| > 1.4 and P-value <0.05 (n=2 replicates). (B) Heat map shows the relative gene expression patterns of transcriptomics datasets performed in (A). **(C)** Bioinformatics analysis of gene expression changes performed with Cancertool to identify down and upregulated pathways in TR cells. **(D)** Jurkat cells were cultured in suspension (control) or embedded in a 3D collagen type I matrix for 3 hours (3D). Cells were collected, lysed, and RNA was extracted for the analysis of transcriptional changes by microarray. Volcano plot shows significant downregulated (blue) and upregulated (red) transcripts in both conditions. |Fold Change| > 1.4 and P-value <0.05 (n=3 replicates). **(E)** Cells were cultured in suspension or embedded in 3D collagen gels for 1h. Then, mRNAs were isolated and the number of differentially expressed genes was analyzed by microarray. Heat map shows the relative gene expression patterns of transcriptomics datasets (n=3). **(F)** Control or TR cells were lysed, mRNA was isolated, and transcriptional changes were analyzed by mRNA expression microarray. **(G)** Graphs show the signal intensity of EU and RNA pol II in control and TR cells in (G). Mean n=84-99 cells ± SD. (5 independent replicates). *** P<0.001. **(H)** Graph shows the correlation between the levels of RNA pol II and the EU incorporation in control and TR cells. n=84-99 (3 independent replicates).

### Persistent 3D cell confinement alters cell cycle and survival alterations

Confined conditions and lamin expression regulate the proliferation and cell cycle progression of cells (35, 36). Thereby, we investigated whether TR cells may present differences in their proliferative ratio. We found that the proliferative ratio of TR cells showed a slight reduction compared to control cells (**Figure 6A**). By analyzing the BrdU (bromodeoxyuridine) incorporation, we found no significant differences in the capacity of TR cells to divide (**Figure 6B**). Furthermore, we analyzed the cell cycle progression of control and TR cells and observed a minor effect of TR cells to accumulate in the G1 phase, without significant increment in the population showing higher DNA amount (**Figure 6C**). To further characterize whether TR cells might show polyploidy, we analyzed the presence of polyploid species in control and TR cells, and confirmed no significant changes in the amount of DNA nor the presence of hyperploidy in TR cells (**Suppl. Figure S6**), indicating that TR cells showed an aberrant nuclear morphology without affecting the polyploidy of the cells. We tested the expression of cyclins and confirmed that TR cells exhibited diminished levels of cyclin A (a typical cyclin for late phases of the cell cycle) compared to control cells (**Suppl. Figure S6**). Cell migration through confined spaces compromises nuclear integrity and promotes DNA damage (37). To determine how DNA damage might be connected to persistent 3D confined conditions, we visualized γH2AX (a well-known DNA damage marker), and found that nuclei of TR cells showed more foci of γH2AX per nucleus than control cells (**Figure 6D, 6E**). We further confirmed these results by measuring the total levels of γH2AX in control and TR cells (**Suppl. Figure S6**), and by quantifying other DNA damage markers, such as phospho-ATM (pATM) (**Suppl. Figure S6**). As we found high levels of this DNA damage marker in TR cells, we characterized their DNA repair response by treating control and TR cells with bleomycin, a drug that induces DNA damage. We visualized the levels of DNA damage by single-cell electrophoresis (comet assay) and found that TR cells were more sensitive to DNA damage than control cells (**Figure 6F, 6G**). Moreover, given the potential implication of the basal levels of DNA damage response on the survival capacity of the cells, we tested whether TR cells were more sensitive to conventional chemotherapies than control cells. Interestingly, we confirmed that TR cells showed more sensitivity to cell death in response to several chemotherapeutic drugs than control cells (**Figure 6H and Suppl. Figure S6**). Together, our results suggest that during persistent 3D confinement, cells undergo permanent changes that alter their proliferative and apoptotic responses.

**Figure 6.**
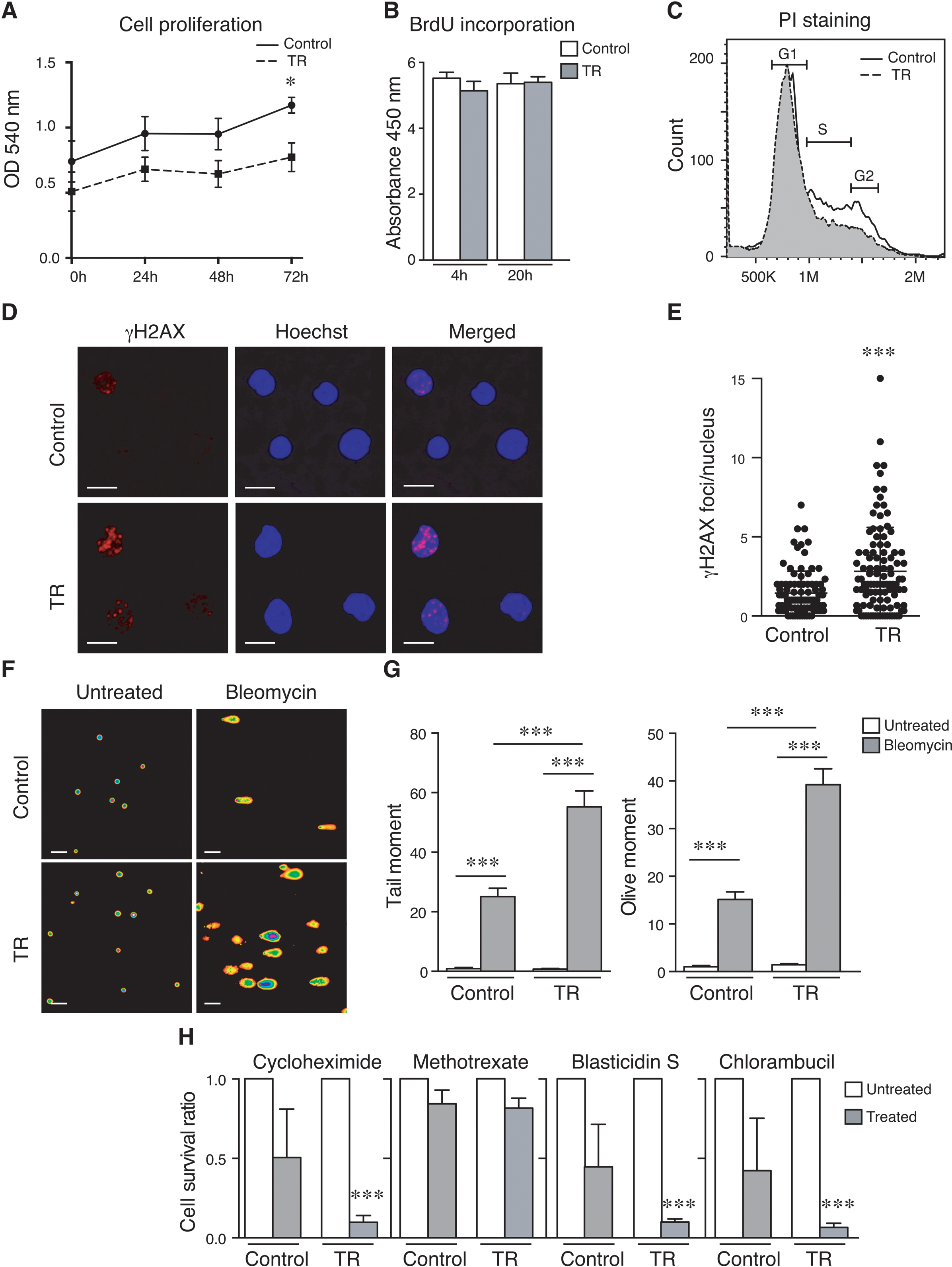
TR cells from persistent 3D confinement show alterations in their cell cycle and survival response. **(A)** Control or TR (dark and dashed lines, respectively) cells were cultured and their cell proliferation was quantified by MTT assay at indicated times. Mean n = 5 replicates ± SD. **(B)** Control and TR cells were incubated with 5-bromo-2’-deoxyuridine (BrdU) for 4 and 20 h. Then, cells were fixed and BrdU incorporation was quantified. Mean n = 3 replicates ± SD. **(C)** Control (white) and TR (grey, dashed line) cells were fixed, permeabilized and stained with propidium iodide. Then, cell cycle progression was analyzed by flow cytometry. Graph shows the G1, S and G2/M phases according to DNA content. **(D)** Control and TR cells were collected, seeded onto poly-L-lysine coated slides, fixed and stained for γH2AX (red) and Hoechst (blue). Bar 10 µm. **(E)** Graph shows the number of foci per nucleus of control or TR cells. n=247-252 (3 independent replicates). **(F)** Control and TR cells were treated with 40 µM Bleomycin for 4 hours, embedded in agarose and lysed. Alkaline comet assay by electrophoresis was performed to visualize the DNA by fluorescence microscopy. Bar 40 µm. **(G)** Graph shows the tail moment analysis of the comet assay in (E). Mean n=302 cells ± SD. **(H)** Control and TR cells were cultured in the presence of the indicated drugs (1µM, Methotrexate; 5 µg/mL, Cycloheximide; 5 µg/mL, Blasticidin S; or 10 µg/mL, Chlorambucil) for 24 h. Then, cells were collected and stained for annexin V-FITC and propidium iodide for flow cytometry analysis. Graph shows the survival ratio of control and TR cells upon normalization to their untreated conditions. Mean n = 3 replicates ± SD. *** P<0.001.

### Cellular changes induced by 3D confinement reduce cell invasiveness

To determine other functions altered in TR cells, we evaluated their migration efficiency. Firstly, we performed a transwell migration assay and found that TR cells showed a defective migration in response to serum (**Figure 7A**). To discard that differences in chemotaxis may be regulated by defective cell adhesion, we evaluated the adhesion of control and TR cells to VCAM-1 (vascular adhesive molecule-1). We did not observe any difference in the levels of TR cells attached to VCAM-1 (**Suppl. Figure S7**). Due to the role of actin polymerization on aberrant lamin B1 distribution, and also the potential impact on the spreading of TR cells during the adhesion process, we tested how control and TR cells spread onto 2D surfaces with non-specific ligands. As expected from the chemotactic assays, we found no differences in the cell adhesion kinetic of control and TR cells nor at short times, and only after 2h a shrinking of the cell area in control cells could be observed (**Figure 7B and 7C**). To further determine the migration of TR cells *in vitro*, we analyzed their capacity to infiltrate into a collagen matrix and found that the deepness invasion of TR cells was not impaired (**Figure 7D and 7E**). Then, we assessed how TR cells might have altered their invasive capacity *in vivo*. For this, we injected an equal number of control and TR cells labeled with vital dyes into the tail vein of immunodeficient recipient mice and analyzed the homing of cells into the bone marrow, liver, and spleen at 24h post-injection. Our analysis indicated a reduction of TR cells reaching these organs compared to controls (**Figure 7F and Suppl. Figure S7**). Together, these results indicate that cell confinement at long-term reduces the capacity of TR cells to invade and reach other tissues.

**Figure 7.**
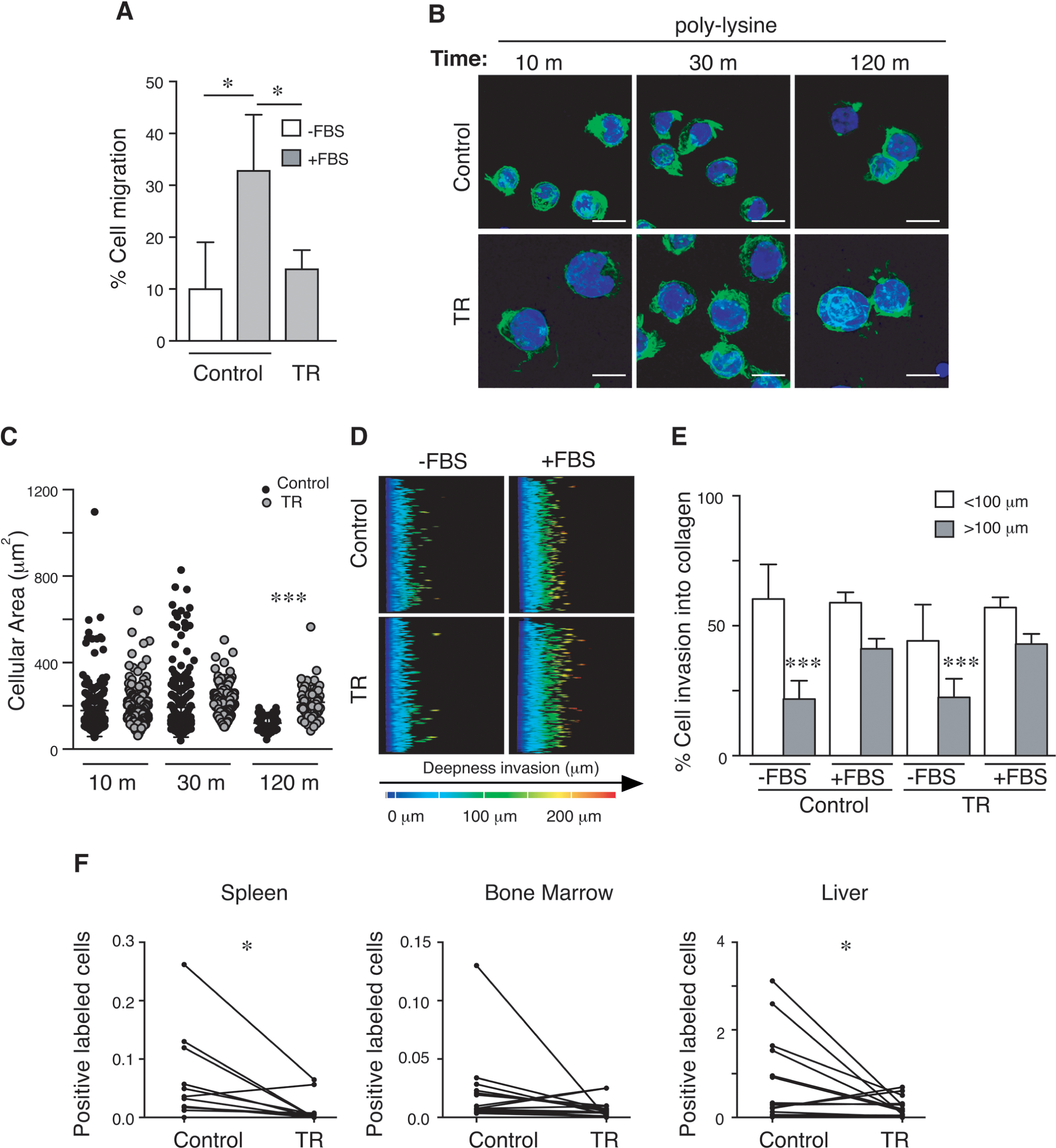
Aberrant cells after suffering 3D confinement exhibit reduced cell invasiveness. **(A)** Control and TR cells were seeded on the top chamber of Transwell inserts and allowed to migrate through 5 µm pores towards FBS for 24 h. Graphs show the percentage of migrated cells. Mean n = 3 replicates ± SD. **(B)** Control and TR cells were seeded on poly-L-lysine coated slides, fixed at the indicated times and stained for with phalloidin (green) and Hoechst (blue). Bar 10 µm. **(C)** Graph shows the cellular area analysis of the adhesion kynetics in (B). Mean n=55-256 cells ± SD (3 independent replicates). **(D)** Control and TR cells were seeded on the top of a collagen 3D matrix and cultured at 37°C for 24 h. Then, cells were fixed, stained with propidium iodide, and serial confocal images of the matrix were taken to calculate the depth invasion of ALL cells towards FBS. **(E)** Graph shows the percentage of cells invading into the collagen gel deeper than 100 µm. Mean n = 6 replicates ± SD. **(F)** Control (Far Red+) and TR (CFSE+) cells were stained with vital dyes, mixed 1:1 and injected into the tail vein of NSG (NOD scid gamma mouse) mice. After 24 h, labeled cells in spleen, bone marrow and liver were determined by flow cytometry. Graph shows the correlation between the control and the TR cells infiltrated into the tissues for each mouse. Mean n=14. * P<0.05. *** P<0.001.

### Persistent 3D confinement confers different biophysical properties and mechanical homogeneity in the nucleus

Nuclear deformability is critical during cell migration through physical barriers such as the endothelium and the interstitial spaces (38–40). Since the nuclear size and morphology of TR cells were affected, we hypothesized that the nuclear deformability would be also affected. To determine how the nucleus deforms upon external forces, we used a cell confiner device, to induce nuclear deformation by global compression. We found that isolated nuclei from TR cells increased their nuclear deformability compared to isolated nuclei from control cells (**Figure 8A, 8B**). Similar changes in the nuclear deformability were observed intact control and TR cells (**Suppl. Figure S8**), indicating that persistent cell confinement indeed may affect the mechanical response of the nucleus. The chromatin compaction also contributes to the mechanical properties of the nucleus (19), and we first interrogated whether TR cells alter their chromatin organization. We assessed DNAse I sensitivity assay to determine the global chromatin compaction of control and TR cells. We found that TR cells exhibited more sensitivity to DNA digestion than control cells, suggesting a more relaxed chromatin compaction (**Figure 8C, 8D**). To further explore this possibility, we treated control and TR cells with trichostatin A (TSA), a chemical drug to induce chromatin decompaction. We revealed that TSA treatment significantly increased the nuclear size of control cells, while TR cells were less sensitive to this treatment, indicative of a lower level of chromatin compaction than control cells (**Suppl. Figure S8**). To investigate these changes in chromatin, we used STORM (Stochastic Optical Reconstruction Microscopy) imaging to observe the spatial organization and density of chromatin within isolated nuclei. We observed a high density of chromatin stained by SiR-DNA around the periphery of the nucleus in control nuclei, whilst TR nuclei displayed an altered chromatin distribution with a decrease in peripheral density and an increase in staining within the nuclear body, as observed with EM (**Suppl. Figure S1**). SIM (Structured Illumination Microscopy) imaging was used to observe the spatial organization of chromatin within whole cells (**Suppl. Figure S9**). These images are consistent with the altered chromatin staining observed using STORM, which confirms that isolation of the nuclei did not alter chromatin organization. Overall, the staining was similar to the distribution of lamin B1 within both cell lines observed in **Figure 2A**, which suggests a potential link between the lamina and chromatin organization, along with accessibility. Osmotic stress has been used to determine the mechanical response of the nucleus (40), as swelling conditions might affect the chromatin compaction and the water diffusivity inside the nucleus. We added KCl or EDTA to isolated nuclei from control and TR cells to determine how the nucleus might respond to swelling conditions. We observed that both treatments increased the area of isolated nuclei from control and TR cells. To address whether actin polymerization might influence on these differences, we treated TR cells with latrunculin and found that these isolated nuclei showed a nuclear swelling more similar to control cells (**Suppl. Figure S9**). We also confirmed that latrunculin treatment reduced the chromatin sensitivity of TR cells to the enzymatic activity of DNAse (**Suppl. Figure S9**).

**Figure 8.**
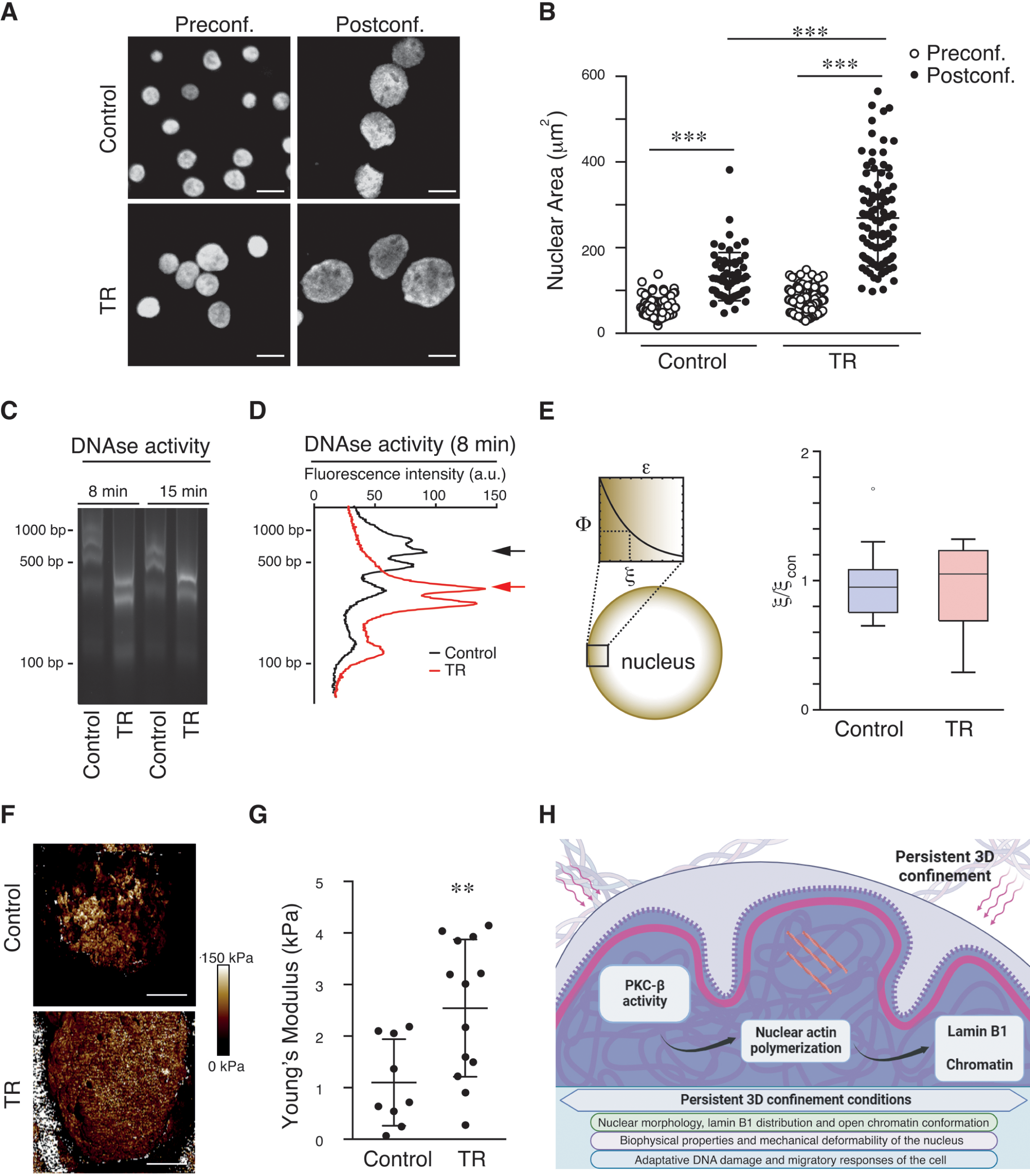
Persistent 3D confinement modulates nuclear biophysics and homogeneity. **(A)** Isolated nuclei from control and TR cells were seeded on poly-L-lysine coated slides, stained with Hoechst, and confined mechanically up to 3 µm. Confocal images pre- and post-confinement are shown. Bar 10 µm. **(B)** Graph shows the quantification of nuclear area pre- and post-confinement from (A). n=66-227 cells ± SD (3 independent replicates). **(C)** Control and TR cells were collected, and their DNA was digested with DNAse I for 8 or 15 min. Then chromatin degradation was resolved in an agarose gel. **(D)** Graph shows the DNA fragmentation profile from control (black) and TR cells (red) as in (D). Arrows indicate the maximum peak of signal in both populations. **(E)** Left panel shows the compaction analysis based on indentation assays of isolated nuclei from control and TR cells performed with optical tweezers. The compaction parameter, Φ, decays exponentially with the indentation depth (shown here in terms of strain, ε, being R the indenter radius). The characteristic length ξ defines the compaction decay. Right graph shows the values for the isolated nuclei from control and TR expressed as the ratio. **(F)** Isolated nuclei from control and TR cells were seeded on poly-L-lysine coated dishes and imaged by AFM (Atomic Force Microscopy). PeakForce Tapping image of a representative nucleus captured with a maximum indentation force of 0.5 nN. **(G)** Graph shows the Young’s modulus values for isolated nuclei from control and TR cells. Each point corresponds to the average value for a nucleus, calculated from >10000 independent force curves. Mean n=9-13 ± SD. **(H)** Schematic representation of the nuclear changes promoted by 3D confinement via PKCβ activity and nuclear actin polymerization. The nucleus alters its morphology, lamin B1 distribution and chromatin compaction. Also, its biophysical properties and mechanical adaptation change compared to control conditions. These alterations also impact on the DNA damage response of the cell and its capacity to invade and infiltrate other tissues *in vivo*. **P< 0.01, *** P<0.001.

Consequently, the aberrant lamin B distribution observed in the TR cells might be linked to alterations on the chromatin organization. We reasoned that the complex set of interactions governing the NE-chromatin crosstalk is encoded as a compaction gradient extending from the nuclear lamina towards the inner nuclear regions (41). Therefore, we hypothesized that this sort of alterations should be also projected as permanent remodeling effects over the isolated nuclear structure. In this sense, we have developed a methodology based on gentle poroelastic indentation using optical tweezers that enables the estimation of the nuclear local density (compaction, Φ) for a certain indentation depth, by merging the information from stiffness and characteristic time of the force relaxation. We applied a range of indentation depths in both isolated nuclei from control and TR cells to reveal the spatial variation of the compaction at the nuclear periphery and found an exponential decay defined by a shape factor of the function, named characteristic length ξ (**Figure 8E**). Although we did not observe significant differences between the average ξ values (**Suppl. Figure S9**), we evinced a noticeable difference in the data dispersion being larger in the TR phenotype (**Figure 8E**), which might indicate that the compaction gradient at the nuclear periphery is more heterogeneous in the TR cells. To further characterize the spatial biomechanical signature of TR cells, we used AFM on isolated nuclei (42). Using PeakForce Capture and PeakForce Tapping, we are able to gain a force curve for each pixel (**Figure 8F**); therefore, we can measure the spatial biomechanical properties. We found that control nuclei are an inhomogeneous material with distinct islands of high stiffness, whilst TR nuclei display a homogeneous stiffness across the nucleus (**Figure 8F**). In this manner, the underlying alterations within the TR cells correlated with a change in the mechanical properties. Calculating the mean stiffness for the nuclei, we could observe an increase in stiffness within the TR nuclei (**Figure 8G**). The increase in stiffness was consistent with a redistribution of lamina and chromatin, which are significant contributing factor to nuclear mechanics. Taken together, our findings indicate that prolonged cell confinement in 3D conditions leads to global changes in the chromatin compaction and biomechanical signature of the nucleus.

## DISCUSSION

Cell migration is a multistep process where migratory cells have to respond to external stimuli to alter their molecular and biophysical properties in order to cross extracellular barriers such as endothelia or the extracellular matrix (7, 26, 43). It is known that the nucleus of migratory cells presents temporary changes during constrained migration. These changes include the mechanical response, nuclear blebs at the leading edge, or even the rupture of the nucleus (17, 18, 44, 45). It was already known that 3D migration controls chromatin changes and alters the transcriptional activity of migratory cells that might persist for days (26, 38); however, how 3D culture conditions promote permanent changes in the nucleus of the cell is not fully understood. In this study, we observed that confined cell culture led to permanent phenotypical changes, including an increment in the size and the number of lobes/invaginations of the nucleus. Aberrant nuclear morphology usually enrolls alterations of the nuclear envelope and chromatin changes that may contribute to cancer heterogeneity and chromatin instability. Therefore, it is possible that 3D constraints might generate physical stress that favors multilobular aberrant nuclear morphologies. Aligning with this, it has been demonstrated that breast carcinoma cells cultured on nanopillars alter their subnuclear morphologies and the distribution of lamin A, concomitant with higher malignancy and cell migration (46). This is particularly important as the lamina network provides the mechanical skeleton to the nucleus, and the levels and the regulation of lamins alter the biomechanical response of the nucleus, the chromatin organization, and the transcriptional activity (2, 5, 38). Phenotypically, our data indicate that cell confinement in 3D induces lamin B1 redistribution in the nucleus of TR cells, showing a similar mechanoresilient adaptation observed in other tumor cells (47). Changes in the localization or distribution of lamin B might be of particular interest, due to lamin-A type levels, but not lamin B, reflect changes in the mechanical properties of the nucleus (48). It has been described that the aberrant nuclear morphology serves as a pathological marker for cancer cells, as well as the number of nucleoli (49). TR cells showed a more aberrant nuclear shape, with significant differences in the intensity of their nucleoli without affecting their number. Furthermore, we have confirmed that lamin B1 interacts with NPM1; however, despite the aberrant distribution of lamin B1, our data suggest that this nuclear redistribution of lamin B1 might occur independently of regulating nucleolar changes or its interaction with nucleolar components. This indicates that the cellular transformation induced by 3D conditions might alter lamin B1 distribution and nuclear morphology, according to malignant transformation controlled by external stress.

Lamin B1 alters its distribution during the cell cycle and might suffer phosphorylation by multiple kinases (50). Consistent with the idea that PKC activity controls lamin B distribution, our findings show that blocking or silencing PKCβ recovers the normal distribution of lamin B at the nuclear periphery. Several PKC isoforms regulate actin polymerization (51, 52), and we found that latrunculin treatment also abrogates the aberrant distribution of lamin B in TR cells. When we interrogated whether latrunculin treatment may regulate lamin redistribution downstream of PKC activity, we found that actin operates connected and downstream of PKC, as the disruption of actin polymerization bypasses the effect of PKC. Interestingly, several PKC isoforms might translocate into the nucleus, including PKCβ (24, 53). Also, in addition to their role in the cytoplasm, the actomyosin machinery (actin, myosins, and associated proteins) can be recruited into the nucleus interacting with polymerases and transcription factors regulating multiple functions, including DNA repair, transcription, and cell cycle progression (54, 55). By using the expression of NLS-ActinR62D, we demonstrated that inhibiting nuclear actin polymerization was sufficient to rescue the normal phenotype of lamin B1, indicating a fundamental role of nuclear actin to control lamin B1 distribution in TR cells. The reasons for these remain unclear and continued studies are needed to better understand specific functions of nuclear actin and PKC isoforms, and how 3D cell culture might contribute to the activation and these molecular processes.

The mechanical stress controls the transcriptional activity of the cell (10, 56). Recent evidence demonstrates that cell growth of human fibroblasts on geometric constraints alters the transcriptional program of the cell and induces cell-fate transitions (33). These changes induced during the lateral confinement in geometric micropatterns are mainly mediated by Lef1 signaling (57). It has been recently proposed that confined migration through narrow spaces might advantage both the selection and induction of phenotypical changes similar to the metastatic process; however, these mechanisms might differ between microfluid devices and 3D matrices (58). Interestingly, we show an increment in the transcriptional activity of TR cells. This aligns with our observation that TR cells increased the signal intensity of nucleolin. Furthermore, our data also indicate higher levels of RNA polymerase II in TR cells, which correlates with higher levels of EU incorporation, reinforcing the idea that TR cells present an upregulation in their transcriptional activity. Interestingly, this transcriptional stress differs from changes at short times of 3D conditions, demonstrating that short- or long periods of mechanical inputs could generate mechanical acute responses or long-term adaptations that might be a factor for genomic instability.

Previous reports have demonstrated that the two major contributors to the DNA repair and DNA damage response (DDR) pathways, ATM and ATR, have been involved in the nuclear mechanics and lamin disposition during cancer cell migration (59, 60). Our functional characterization of TR cells showed a reduction in their invasion capacity in vivo, and more DNA damage markers signal in resting conditions, suggesting that these cells might suffer higher genomic stress and a predisposition to become more sensitive to specific treatments. This might suggest that nuclear changes induced by 3D conditions might have a significant impact on cancer cell invasion and genome heterogeneity. On the other hand, our observations also align with previous work demonstrating that nuclear actin polymerization controls cellular proliferation and migration (61), suggesting that nuclear actin homeostasis regulated by 3D conditions might present additional roles in controlling functional changes observed in TR cells.

Preventing chromatin changes resulted in impaired migration of multiple cell types through confined conditions (40, 41). We found that TR cells showed less compacted chromatin, which results in more sensitive nucleosomes to DNAse activity, and redistributed around the nucleus in a similar manner to the redistribution of lamin B. In agreement with these studies and our observations using DNAse, TR cells showed a minor nuclear expansion under TSA treatment, indicating a lower chromatin compaction *per se*. It is plausible that the less condensed chromatin of TR cells might be a key factor to prime these cells for different nuclear functions. This might be particularly important as chromatin changes observed in microfluidic devices might not operate in 3D collagen matrices (39). As both, the chromatin and the nuclear lamina disposition seemed to be altered in TR cells, we assessed how 3D confined conditions might promote persistent changes in the mechanical signature of the nucleus. First, we observed by optical tweezers that the abnormal lamin distribution found in TR cells might be linked to a more heterogeneous chromatin compaction at the nuclear periphery. This heterogeneity might be due to the force balance at the nuclear edge. In general terms, the spheroidal shape of the cell nucleus is defined by the mechanical balance between the osmotic pressure (outwards) and the Laplace pressure (inwards), which is defined by the mechanical stress at the interface (62). As a consequence, it is expected that a more compact and homogeneous periphery would define a nuclear structure without geometrical heterogeneities, as observed in the control nuclei. Conversely, the existence of regions with a weakened surface tension can rise to the formation of mechanical instabilities as those observed in nuclear blebs or in laminopathies (63). When we used a specific AFM approach to identify the continuum biophysical signature of isolated nuclei, we showed that aberrant nuclei from TR cells present an increased stiffness, which correlates with morphological alterations and changes in the DNA susceptibility to enzymatic degradation. It has been reported how AFM and another mechanical techniques differ considerably in the quantified response (64). Our results suggest that the differences between the nuclear compression, AFM and optical tweezers might be likely due to more complex factors contributing to the mechanical response of the nucleus, including a higher level of chromatin decondensation, diminution of actin polymerization, and loosening of the nuclear lamina, which is an interesting difference that will need further studies in the future.

Overall, our findings describe how 3D cell culture conditions alter the nuclear morphology, chromatin conformation, lamin B1 distribution, and the biomechanical response of the nucleus (**Figure 8H**). Considering the functional changes observed in gene expression, the adaptative cell survival to drugs, and the migration of TR cells; our observations may contribute to explaining novel biomechanical effects of 3D conditions in the nuclear biology that might occur during cancer cell invasion, tissue repair, and developmental processes.

## DATA AVAILABILITY

Microarray data have been deposited with the Gene Ontology Database under accession numbers GSE181375 and GSE226621. All relevant data are available from the corresponding author upon reasonable request.

## ETHICS APPROVAL

All procedures for animal experiments were approved by the Committee on the Use and Care of Animals and carried out in strict accordance with the institution guidelines and the European and Spanish legislations for laboratory animal care.

## SUPPLEMENTARY DATA

Supplementary Data are available at NAR online.

## AUTHOR CONTRIBUTIONS

R.G.N., H.Z.C., conducted experiments and contributed to data interpretation and discussion. A.D.L., conducted experiments and contributed to data interpretation. H.L.M. conducted experiments and contributed to data interpretation. P.R.N., F.M. supervised the experiments related to the optical tweezers, contributed to data interpretation and provided financing. L.W., designed and supervised the experiments related to STORM. C.P.T., designed and supervised the experiments related to STORM and AFM, contributed to data interpretation, and provided financing. J.R.M designed and supervised the experiments, contributed to data interpretation, wrote the paper, and provided financing.

## ACKNOWLEDGEMENT

The authors thank the Microscopy Unit of Instituto de Investigación Biosanitaria Gregorio Marañón (IiSGM) for assistance with confocal analyses. The authors are also grateful to the EM and Animal Facilities of platforms of the CIB Margarita Salas for their assistance and support with the EM and in vivo experiments. The UCM-Genomic CAI Unit for their assistance with microarray experiments. We thank Robert Moorhead and Daniel. E. Rollins for assistance and support at Royce@Sheffield. We thank Tobías Álvaro-Thomsen for assistance and his support by Asociación Española Contra el Cáncer.

## FUNDING

We wish to acknowledge the Henry Royce Institute for Advanced Materials, funded through Engineering and Physical Sciences Research Council (EP/R00661X/1, EP/S019367/1, EP/P02470X/1 and EP/P025285/1). This research was supported by a FPI Scholarship 2018 (Ministerio de Ciencia e Innovación/MICINN, Agencia Estatal de Investigación/AEI y Fondo Europeo de Desarrollo Regional/FEDER) to R.G.N.; grants from Comunidad de Madrid (Y2018/BIO-5207) and from the Ministerio de Ciencia e Innovación (MICINN) Agencia Estatal de Investigación (AEI) (PID2020-115444GB-I00, /AEI/10.13039/501100011033) to P.R.N; grants from the Ministerio de Ciencia e Innovación (MICINN) Agencia Estatal de Investigación (AEI) (PID2019-108391RB-100), and Comunidad de Madrid (Y2018/BIO-5207, S2018/NMT-4389 and REACT-EU program PR38-21-28 ANTICIPA-CM) to F.M.; Medical Research Council (MR/M020606/1) and Science and Technology Facilities Council (19130001) to C.P.T.; and grants from 2020 Leonardo Grant for Researchers and Cultural Creators (BBVA Foundation), Ayuda de contratación de ayudante de investigación PEJ-2020-AI/BMD-19152 (Comunidad de Madrid), and the Ministerio de Ciencia e Innovación (MICINN) Agencia Estatal de Investigación (AEI) (PID2020-118525RB-I00, AEI/10.13039/501100011033) to J.R.M.

## CONFLICT OF INTEREST

Authors declare no conflict of interest.

## Notes

### Competing Interest Statement

The authors have declared no competing interest.

### Summary of Updates

New Figures, discussion and supplemental material

