## Supplemental Info for "3D environment promotes persistent changes in lamin B1 distribution, the biomechanical signature of the nucleus, and adaptative survival and migratory functions"

**Supplementary Materials and Methods.**

**Supplementary Tables S1-S3.**

**Supplementary Videos S1-S4.**

**Supplementary Figures S1-S8.**

### SUPPLEMENTARY MATERIALS AND METHODS

#### Antibodies and cellular dyes

| Antibodies | Conc. of use | Host | Company | #Cat |
|---|---|---|---|---|
| Anti-laminB1 | 1/100 | Mouse | Santa Cruz | 365962 |
| Anti-β-Actin | 1/1000 | Rabbit | abcam | ab227387 |
| Anti-PKC-β | 1/1000 | Rabbit | Thermo Fisher | PA5-13740 |
| Anti-pH2AX | 1/100 | Mouse | Invitrogen | 14-9865-82 |
| Anti-cyclin A | 1/100 | Rabbit | EMD Millipore | 06-138 |
| Anti-pATM | 1/100 | Mouse | Invitrogen | MA1-2020 |
| Anti Nucleolin | 1/100 | Mouse | Sigma-Aldrich | N2662 |
| Anti-NPM1 | 1/1000 | Mouse | Cell Signaling | 3542S |
| Anti RNA polymerase II (4H8) | 1/100 | Mouse | Novus | NB-200-598 |
| Click-iT™ RNA Alexa Fluor™ 594 | 1 mM 1h | - | Thermo Fisher | C10330 |
| siR DNA | 1/1000 | - | Spirochrome | sc007 |
| CF 488A anti-mouse IgG | 1/200 | Donkey | Biotium | 20014 |
| CF647 anti-rabbit IgG | 1/200 | Goat | Biotium | 20045 |
| Hoechst 33342 | 1/1000 | - | Invitrogen | H3570 |
| Popidium iodide | 1/10000 | - | Sigma-Aldrich | P4864 |
| CellTrace™ Far Red | 2 µM 30 min | - | Invitrogen | C34564 |
| CellTrace™ CFSE | 5 µM 30 min | - | Invitrogen | C34554 |
| Invitrogen™ BCECF, AM | 1 µM 30 min | - | Invitrogen | 10646693 |

#### Drug treatments and transfections

| Drug | Conditions of use | Company | #Cat |
|---|---|---|---|
| Latrunculin B | 2 µg/mL 1 h | Enzo | BMC-T110_0001 |
| Jasplakinolide | 1 µg/mL 1 h | Enzo | ALX-350-275 |
| Enzastaurine | 2 nM 30 min | Sigma-Aldrich | C050 |
| Staurosporine | 50 nM 30 min | Sigma-Aldrich | 19-123-M |
| PMA | 50 ng/mL 15 min | Selleckchem | S7791 |
| Bleomycin | 40 µM 4 h | Alfa Aesar | J60727 |
| Cycloheximide | 5 µg/mL 24 h | Sigma-Aldrich | C-6255 |
| Chlorambucil | 10 µg/mL 24 h | Sigma-Aldrich | C-0253 |
| Methotrexate | 1 µM 24 h | Sigma-Aldrich | M9929 |
| Blasticidin S | 5 µg/mL 24 h | Calbiochem | 203350 |
| Trichostatin A | 300 nM 18 h | Cell Guidance Systems | SM36-1 |
| EDTA | 5 mM 10 min | Panreac | A1103,0250 |
| KCl | 5 mM 10 min | Sigma-Aldrich | P-4504 |

**Cell transfection.** For PKCβ silencing, TR cells were transfected with siTRANS siRNA transfection reagent (Origene Technologies) in RPMI following the manufacturer's instructions and with specific pool of three to five target-specific 19-25 nt siRNAs designed to knockdown PKCβ (sc-29450) or control siRNAs (Santa Cruz Biotechnology). After 24 hours, the transfection efficiency was confirmed by western blotting and cells used for the indicated experiments. For transient expression of actin mutants in the nucleus, 500 ng of NLS-YFP-WT (wild type actin), NLS-YFP-R62D (R actin) and NLS-YFP-S14C (S actin) DNA plasmids were used as in (65). In brief, cells were transfected using Lipofectamine 2000 (Invitrogen). After 48 h, cells were seeded on poly-L-lysine coated slides, fixed and subjected to indirect immunofluorescence.

**Western blotting.** Cells were washed with cold PBS, lysed in Laemmli buffer (BioRad) and then sonicated (15 s at 70% amp) using a Microson XL2000 (Misonix). Protein samples were boiled, resolved by electrophoresis in 7.5-15% acrylamide gels and transferred to nitrocellulose membranes (GE Healthcare Life Science). Blocking was performed with 3% low fat milk in TBS-Tween (0.1%) for 1 h at room temperature (RT) and incubated overnight with primary antibodies (1:1000) at 4ºC with rotation. After washing with TBS-Tween 0.1% membranes were incubated with HRP-labelled secondary antibodies (1:5000) for 1 h at RT. Protein signal was developed using the enhanced chemiluminescent detection method (Amersham) and analyzed in a ChemiDoc (Bio-Rad). For the stripping process, 3 stripping buffer washes (1% Glycine, 1% SDS and 0.05% NP-40 pH=2.2) of 30 min were performed before blocking the membranes and reprobing with specific antibodies.

**Flow cytometry.** Isolated nuclei were resuspended in TKMC buffer and nuclear size and complexity were measured with a CytoFlex (Beckman Coulter) as described in (64). For intracellular proteins quantification, cells were fixed with PFA 4% for 5 min at RT, permeabilized with PBS Tx-100 0.1% for 5 min, blocked with human IgG (1:1000 in PBS; 30 min) and incubated with specific primary antibodies of interest (1:100 in PBS) overnight at 4ºC. Cells were washed with PBS 3 times and incubated with the appropriated fluorochrome-conjugated secondary antibody (1:100) for 30 min. To analyze the cell cycle progression, $3x10^5$ cells were fixed in ice-cold 70% (v/v) ethanol overnight at -20ºC. Then, cells were washed with PBS and incubated in PBS with RNAse (100 µg/ml), 0.1% Tx-100 and propidium iodide (50 µg/ml) for 30 min at RT. Cells were washed, resuspended in PBS and G1, S, and G2/M phases were determined by flow cytometry. Ploidy was determined by DNA content detection upon Hoechst 33342 (10 µg/mL, 2 hours) staining of control and TR cells. All data were analyzed using the Flow Jo software (Flow Jo LLC, Ashland, OR, USA).

**Electron microscopy.** $3x10^6$ cells were fixed for 1h in 3% glutaraldehyde in PBS and then washed twice with PBS. After post-fixation in 1% osmium tetroxide and 0.8% potassium ferricyanide for 1h at 4ºC, samples were washed with PBS prior to dehydration with an increasing gradient of ethanol (30%, 50%, 70%, 80%, 90% and 100%) of 10 min per step. Then, samples were embedded in LX112 resin and polymerized for 48 h at 60ºC. 60-80 nm sections were placed in copper grids of 75 mesh and stained with 5% uranyl acetate for 30 min and lead citrate for 4 min. Samples were viewed in a JEOL 1230 TEM and images were taken with a CMOS TVIPS 16 mp camera. Nuclear shape was analysed by segmentation with ImageJ.

**Stochastic Optical Reconstruction Microscopy (STORM) Imaging.** For sample preparation and imaging, we followed the protocols in (20, 65, 66). Briefly, pre-cleaned No 1.5, 25-mm round glass coverslips were placed in 6-well cell culture dishes and coated with 100 µg/mL poly-D-lysine. $6x10^6$ isolated nuclei from control and TR cells were seeded on the coverslips, allowed to attach for 2 h at 37ºC, and fixed in pre-warmed 4% (w/v) PFA in PBS. Residual PFA was quenched for 15 min with 50 mM ammonium chloride in PBS. After 3 TBS washes, nuclei were subjected to permeabilization with cold TBS-Tx-100 0.5% for 10 min, washed with TBS, and blocked with 3% BSA, 0.1% Tx-100 in TBS for 1 h. Then, nuclei were incubated with 1 µg/mL SiR-DNA (Spirochrome) for 1 h and washed in TBS. After 15 min post-fixation with 4% PFA, nuclei were stored at 4°C in the dark, in 0.02% NaN3 in PBS. Before STORM imaging, coverslips were assembled into the Attofluor® cell chambers (Invitrogen). Imaging was performed in freshly made STORM buffer consisting of 10% (w/v) glucose, 10 mM NaCl, 50 mM Tris–pH 8.0, supplemented with 0.1% (v/v) 2-mercaptoethanol and 0.1% (v/v) pre-made GLOX solution (5.6% (w/v) glucose oxidase and 3.4 mg/ml catalase in 50 mM NaCl, 10 mM Tris–pH 8.0), which was stored at 4 °C for up to a week.

Imaging was undertaken using the Zeiss Elyra PS.1 microscope. Illumination was from a HR Diode 642 nm (150 mW) laser delivering 7–14 kW/cm2 power density on the samples through a 100× NA 1.46 oil immersion objective lens (Zeiss alpha Plan-Apochromat). Imaging was performed under highly inclined and laminated optical (HILO) illumination to reduce the background fluorescence. The final image was projected on an Andor iXon EMCCD camera after passing a BP 420–480/BP495–550/LP 650 emission filter. The images were taken with 20 msec exposure per frame for 60 000 frames.

The images were then processed through our STORM analysis pipeline using the Zeiss Zen Black software. Single molecule detection and localization were performed using a 9-pixel mask with a signal-to-noise-ratio of 6 in the 'Peakfinder' settings while applying the 'Account for overlap' function. This function allows multi-object fitting to localize molecules within a dense environment. Molecules were then localized by fitting to a 2D Gaussian function. The image processing was then subjected to model-based cross-correlation drift correction. The final reconstruction was then rendered at 10 nm/pixel and displayed in Gauss mode where each localization is presented as a 2D gaussian distribution with a standard deviation based on its precision.

**Structured Illumination Microscopy (SIM) Imaging.** 2x$10^6$ control and TR cells were seeded on 100 µg/mL poly-D-lysine coated coverslips, allowed to attach for 30 min at 37ºC, fixed and stained with 1 µg/mL SiR-DNA (Spirochrome) following the same protocol as for STORM imaging. Imaging was undertaken using a Zeiss Elyra PS 1 microscope with a 63x NA 1.46 oil immersion objective lens (Zeiss α Plan-Apochromat). A HR Diode 642 nm (150 mW) laser was used to illuminate SiR-DNA and the fluorescence was filtered using a BP 420–480/BP495–550/LP 650 emission filter. The images were taken with 100 msec exposure per frame. The total of 5 rotations of the illumination pattern were implemented to obtain homogeneously weighted two-dimensional information, and the illumination pattern undergone 5 phase stepping in each orientation, leading to a total of 25 raw images for one SIM reconstruction. Super-resolution SIM image processing was performed using the Zeiss Zen Black software.

**Cell adhesion.** Control and TR cells were labeled with 1 µM BCECF-AM (2',7'-Bis-(2-Carboxyethyl)-5-(and-6)-Carboxyfluorescein, Acetoxymethyl Ester) for 30 min in RPMI. Then, cells were washed in complete medium and seeded onto 2.5- 5 µg/mL VCAM-1 coated wells. After 30 min, wells were washed with RPMI and attached cells were lysed in PBS-01% SDS and fluorescence at 490 nm was quantified in a Varioskan (Thermo Fisher). Percentage of adhered cells from the total was calculated.

**RNA Microarray and quantitative real-time PCR (qRT-PCR).** For RNA isolation, $10^6$ cells were lysed using TRI reagent (Sigma) for 30 min. Then, chloroform was added, mixed and incubated for 30 min. After 15 min centrifugation at 13000 rpm, the soluble phase containing the RNA was collected and precipitated with isopropanol for 15 min on ice. The pellet obtained by 15 min 13000 rpm centrifugation at 4ºC, was washed with cold ethanol 80%, centrifugated at 8000 rpm for 5 min and allowed to dry. Total RNA was resuspended in RNAse-free water, and purity and concentration were determined by NanoDrop ND-1000 Spectrophotometer (Fisher Scientific). 1 mg of isolated RNA from each sample was used for microarray analysis by Human Gene Clariom S Assay (Thermo Fisher Scientific). Data were processed, normalized and log2 transformed by the UCM-Genomic CAI Unit. Analysis was performed using Transcriptome Analysis Console and CancerTool visualization interphase for cancer from cBiogune (66). For Quantitative real-time PCR (qRT-PCR) analysis, 1 mg of purified RNA was retrotranscribed using First Strand cDNA synthesis kit (Thermofisher) and cDNA concentrations were quantified by NanoDrop. Oligonucleotides for selected genes were designed according to the PrimerQuest Tool from IDT: *EHMT2*: 5′-CATCAAGGTGCGGCTACAG-3′ and 5′-ATCCTCTCTCACATCAGCCT-3′; *MEN1*: 5’-GTGAGCTGGTGAAGAAGGTC-3’ and 5’-CACCGGAGCTGTCCAAT-3; and *qTBP*: 5′-CGGCTGTTTAACTTCGCTTC-3′ and 5′-CACACGCCAAGAAACAGTGA-3′. Quantitative real-time PCR (qRT-PCR) was performed using iQ SYBR Green Supermix (Bio-Rad Laboratories) and LightCycler 480 II (Roche). Assays were made in triplicates and results normalized according to the expression levels of TBP. Melt curve analysis was performed at the end of PCR to confirm the presence of a single, specific product. The results were expressed using the ΔΔCt method for quantification.

**EU incorporation assay.** In situ nascent RNA detection was performed with the Click-iT RNA Alexa Fluor 594 Imaging Kit (Invitrogen) following the manufacturer's instructions. In brief, $10^5$ cells were incubated with 1 mM EU (5-etiniluridine) in culture medium for 1 hour. Samples were washed in PBS, fixed in 4% PFA for 5 min and permeabilized with 0.5% Triton-X-100 in PBS for 10 min. Then, samples were incubated with working solution of Click-iT reaction buffer with $CuSO_4$ and Alexa Fluor azide for 30 min at RT, washed and resuspended in PBS. EU incorporation was detected by Alexa Fluor 594 detection by flow cytometry. For confocal imaging detection, $10^5$ cells were sedimented onto 10 μg/mL poly-L-lysine-coated glass slides for 30 min prior to fixation, permeabilization and proceeding as explained above.

**MTT cell proliferation assay.** $10^5$ cells were seeded on a microwell plate in 100 μL of culture medium for 0, 24, 48 or 72 h. Then, 50 μg of the MTT (3-(4,5-Dimethylthiazol-2-yl) labelling reagent (Sigma) was added and cells were incubated for 4 h at 37°C in a humidified atmosphere. MTT was solubilized using 1 volume of isopropyl alcohol and 1/5 volumes of PBS-3% SDS. Absorbance was measured at 560 nm in a Varioskan (Thermo Fisher).

**BrdU cell proliferation assay.** $2x10^4$ cells seeded on a microwell plate with culture medium and the BrdU cell Proliferation Assay Kit (Cell Signaling) was used for cell proliferation quantification following the manufacturer's instructions. Briefly, BrdU was added, and cells were incubated for either 4 h or 20 h at 37ºC to allow them to replicate. Cells were fixed and stained, first with a primary antibody against BrdU for 1h at RT and then with an HRP-conjugated secondary antibody for 30 min at RT. Finally, HRP substrate TMB was added and the cell incorporation of BrdU was measured by the absorbance at 450 nm in a Varioskan.

**Comet assay.** To detect small amounts of DNA damage, including single and double-stranded breaks, the comet assay kit (Abcam) was used following the manufacturer's instructions. Briefly, $10^5$ control and TR cells were treated or not with 40 μM Bleomycin

(Alfa Aesar) for 4 hours at 37°C, washed in cold PBS and combined with agarose (at 37°C) at a ratio of 1:10 in comet slides. The slides were immersed in 4°C lysis buffer for 90 min and later in freshly prepared alkaline unwinding solution for 90 min at 4°C. The slides were then placed in an electrophoresis slide tray with the alkaline solution and electrophoresis was performed (18V for 30 min). The cells in agarose were washed in distilled water, fixed in 70% cold ethanol, dried and stained with 0.1 μg/mL SYBR-Gold for 30 min at room temperature and then viewed. The tail moment (TM=tail length x % of DNA in the tail) and olive tail moment (OTM = (Tail mean – Head mean) × (Tail %DNA)/100) are two common descriptors of DNA damage for the alkaline comet assay, which were analyzed by the CometScore software (TriTek). At least 50 cells were analyzed per sample.

**Cell apoptosis.** Cells were treated with or without the following apoptosis inductors at the indicated concentrations for 24 hours: 1μM, Methotrexate; 5 μg/mL, Cycloheximide; 5 μg/mL, Blasticidin S; or 10 μg/mL, Chlorambucil. Then, cells were collected, washed in PBS and stained with AnnexinV and propidium iodide, according to manufacturer's instructions (Immunostep). The proportion of Annexin V-FITC and propidium iodide negative cells was determined by flow cytometry (CytoFlex, Beckman Coulter) as living cells. Data analysis was performed using the software FlowJo.

**Cell migration through Transwells.** $2x10^5$ cells were resuspended in serum-free RPMI medium and seeded on the upper chamber of a Transwell insert inserts (Corning Costar, 6.5 mm diameter, 5 μm pore size). RPMI supplemented with serum was added to the bottom chamber of the Transwell as chemoattractant. After 24h, migrated cells were collected from the bottom chamber and counted to calculate migration index.

**Cell invasion into 3D gels.** 100 μL collagen matrix was reconstituted at 1.7 mg/mL in RPMI, neutralized with 7.5% $NaHCO_3$ and 25 μM HEPES inside Transwell inserts (0.4 μm, Costar). After 1 h at 37ºC, 100 μL of serum-free RPMI containing $3x10^5$ cells were added on the top of the gel. FBS supplemented RPMI medium was added to the bottom chamber of the Transwell as chemoattractant. After 24 h, embedded cells in the collagen gel were fixed with 4% PFA for 1 h, permeabilized with 0.5% Triton-X-100 in PBS for 30 min and stained with propidium iodide. Invading cells were imaged with a sCMOS Orca-Flash 4.0LT camera (Hamamatsu) coupled 5 to an inverted DMi8 microscope (Leica), capturing serial z- stacks every 10 μm with a 10× objective (dry ACS APO 10x/NA 0.3). 3D reconstructions were performed with Leica software. Percentage of penetrating cells in each range of distance was quantified with ImageJ software.

**Chromatin decompaction.** Control and TR cells were treated or not with 300 nM Trichostatin A (Cell Guidance Systems) for 18 h to allow chromatin decompaction. Cells were seeded on poly-D-Lysine coated glass coverslips for 1 h before 4% PFA fixation for 5 min. Cells were permeabilized with 0.5%-Tx-100 in PBS for 5 min, stained with Hoechst 33342 (1μg/mL) and observed in the microscope.

**DNAse I-sensitivity assay.** $3x10^6$ cells were resuspended in Lysis buffer (Tris-HCl 10 μM, sucrose 300 mM, NaCl 15 μM, $MgCl_2$ 5 μM, NP-40 0.5%, DTT 0.5 μM) with protease inhibitors. Then, lysates were digested with 1 U of DNAse I (Thermo Fisher) diluted in DNAse buffer (Tris-HCl 10 μM [pH 7.5], $MgCl_2$ 2.5 μM, $CaCl_2$ 0.5 μM) for 8 or 15 min at 25°C. Reactions were stopped by adding STOP buffer (Tris-HCl 10 μM, EDTA 5 μM, NaCl 200 μM, SDS 0.2%), and samples were subsequently incubated with 1 μg RNAse A (Sigma) for 30 min at 37°C and 1 μg Proteinase K (Sigma) 2 h at 52°C. Digested DNA was isolated with phenol/chloroform (Thermo Fisher) and precipitated with 2.5 volumes of ethanol and 3 M sodium acetate at −20°C overnight. DNA pellet was dissolved in water and visualized in 1.5% agarose gel with SYBR green (Sigma) in a ChemiDoc (Bio Rad).

**Optical tweezers.** Isolated cell nuclei resuspended in TKM buffer were mixed with polystyrene beads with a mean particle size of 3.0 μm (Sigma-Aldrich) at final concentrations of $10^7$ nuclei/ml and 0.005% (w/v), respectively. The optical tweezers device (SensoCell, Impetux Optics S.L., Spain) is equipped with an ultra-stable single-frequency laser source (5 W, $\lambda$ = 1064 nm) guided by acousto-optic deflectors and includes a direct force measurement platform capable of detecting the light scattered from the optical traps. The OT device is mounted on an inverted microscope (Eclipse Ti, Nikon, Japan) and a water immersion objective (Plan Apo VC 60XA/1.20 WI, Nikon) employed to focus the laser on the sample. A short-pass dichroic mirror transmits the bright-field illumination and reflects the IR trapping beam, and a short-pass filter was used to avoid IR laser radiation leaking. Sample bright-field imaging was captured by a CMOS camera (DCC1545M-GL, ThorLabs, USA) and the optical traps were operated with LightAce software (Impetux Optics S.L.). For each indentation experiment, a volume of 40 μL of the sample was placed in a custom-made glass chamber. This chamber was mounted on the microscope and measurements were performed at 25 ºC. Only average-sized, round, symmetric nuclei with no alterations or major perturbations of their integrity were selected to perform the indentation routine. Indented nuclei got attached to the bottom surface of the glass chamber by unspecific interactions, and indenter beads were placed next to the nuclear envelope at axial positions and approximately 2 μm above the bottom surface of the chamber. Indenter beads had a nominal diameter of 3 μm, but the exact diameter of each bead used for indentation was measured by image analysis using the ImageJ software. The indentation process consisted on pushing the nucleus laterally by generating an oscillation of the indenting bead. The fixed parameters of the oscillation are the shape (squared), frequency (0.5 Hz, which is enough to permit a complete relaxation between consecutive cycles), and offset (100%). The amplitude was variable, and the routine was set to sweep the 0.6-1.6 μm range with steps of 0.05 or 0.10 μm. Data were acquired during 45 s for each amplitude step. For each indentation experiment, the elasticity constant of the trap was calculated by using a particle scan routine included in the LightAce software. We wrote a specific Matlab (MathWorks, USA) script to analyze the data from the files obtained by the custom-made Labview indentation routine. This script is able to isolate each force-time curve for every indentation cycle and to calculate the stiffness from the indentation force values and the water diffusivity from the poroelastic relaxation times.

**Statistics.**

Statistical analysis and comparisons were made with GraphPad Prism6. The numerical data are presented as mean ± SD. Differences between means were tested by Student t test for two groups comparison. Where 3 or more groups were analyzed, one-way ANOVA was performed. P-values are indicated by asterisks ((*) $P < 0.05$; (**) $P < 0.01$; (***) $P < 0.001$).

### SUPPLEMENTARY TABLES

**Supplementary Table S1.** Transcriptional changes of control and TR cells by microarray analysis. |Fold Change| > 1.4 and P-value <0.05. n=2.

**Supplementary Table S2.** Transcriptional changes of cells cultured in suspension or in embedded in a 3D collagen matrix by microarray analysis. |Fold Change| > 1.4 and P-value <0.05. n=3.

**Supplementary Table S3.** Informatics analysis of common transcriptional changes from TR cells and cells embedded in a 3D collagen matrix.

### SUPPLEMENTARY VIDEOS

**Supplementary Videos S1 and S2.** 3D reconstructions from confocal sections of representative lamin B1-stained nuclei from control (S1) and TR (S2) cells.

**Supplementary Videos S3 and S4.** 3D reconstructions from confocal sections of representative emerin-stained nuclei from control (S3) and TR (S4) cells.

**SUPPLEMENTARY FIGURES**

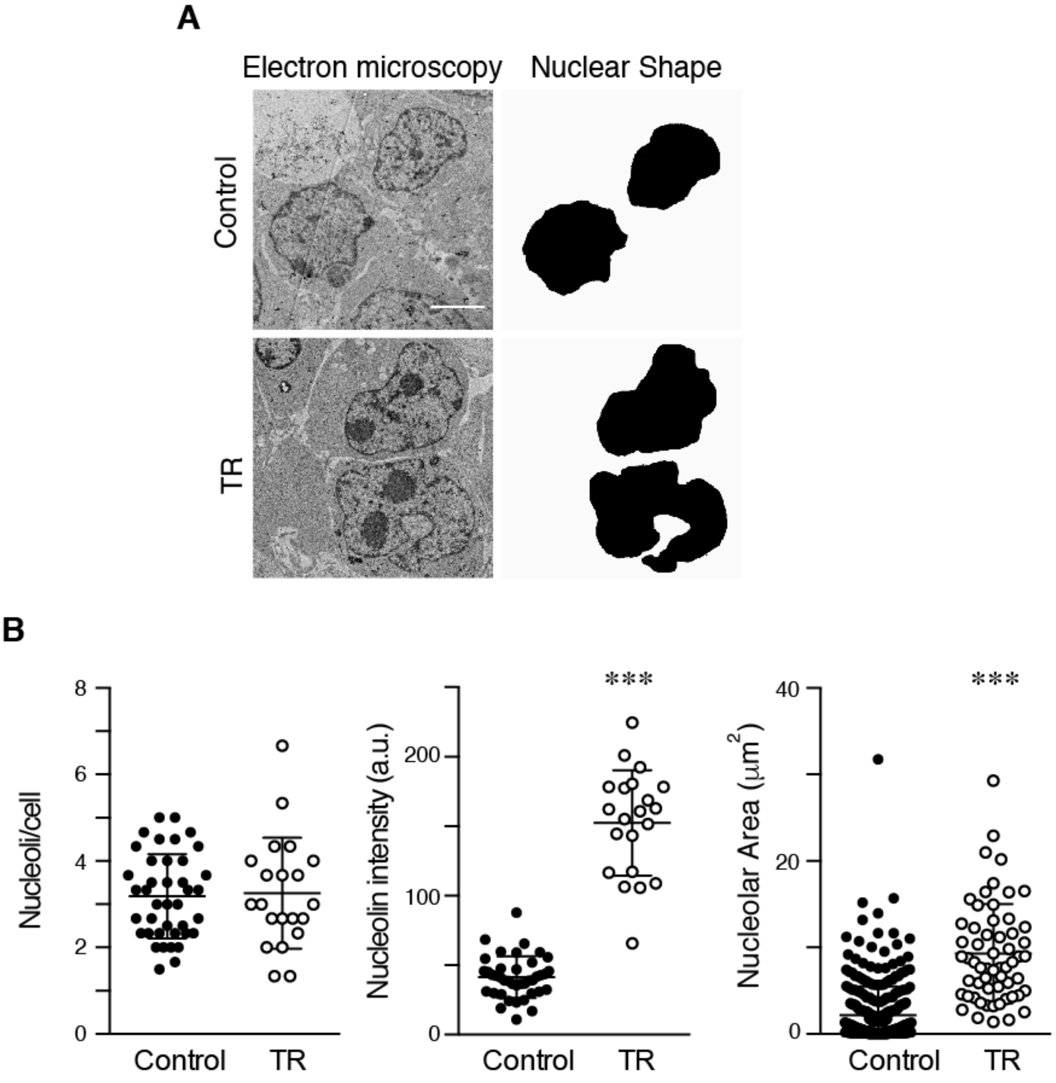


**Figure S1**

**Fig. S1. (A)** Control and TR cells were collected and processed for thin section electron microscopy to visualize the nuclear morphology. Right panels indicate the shape of the nuclei in black. **(B)** Graphs show the number of nucleoli in control and TR cells (left panel), the mean intensity signal of nucleolin (center panel), and the nucleolar area occupied (right panel). Bar 4 μm. *** $P < 0.001$.

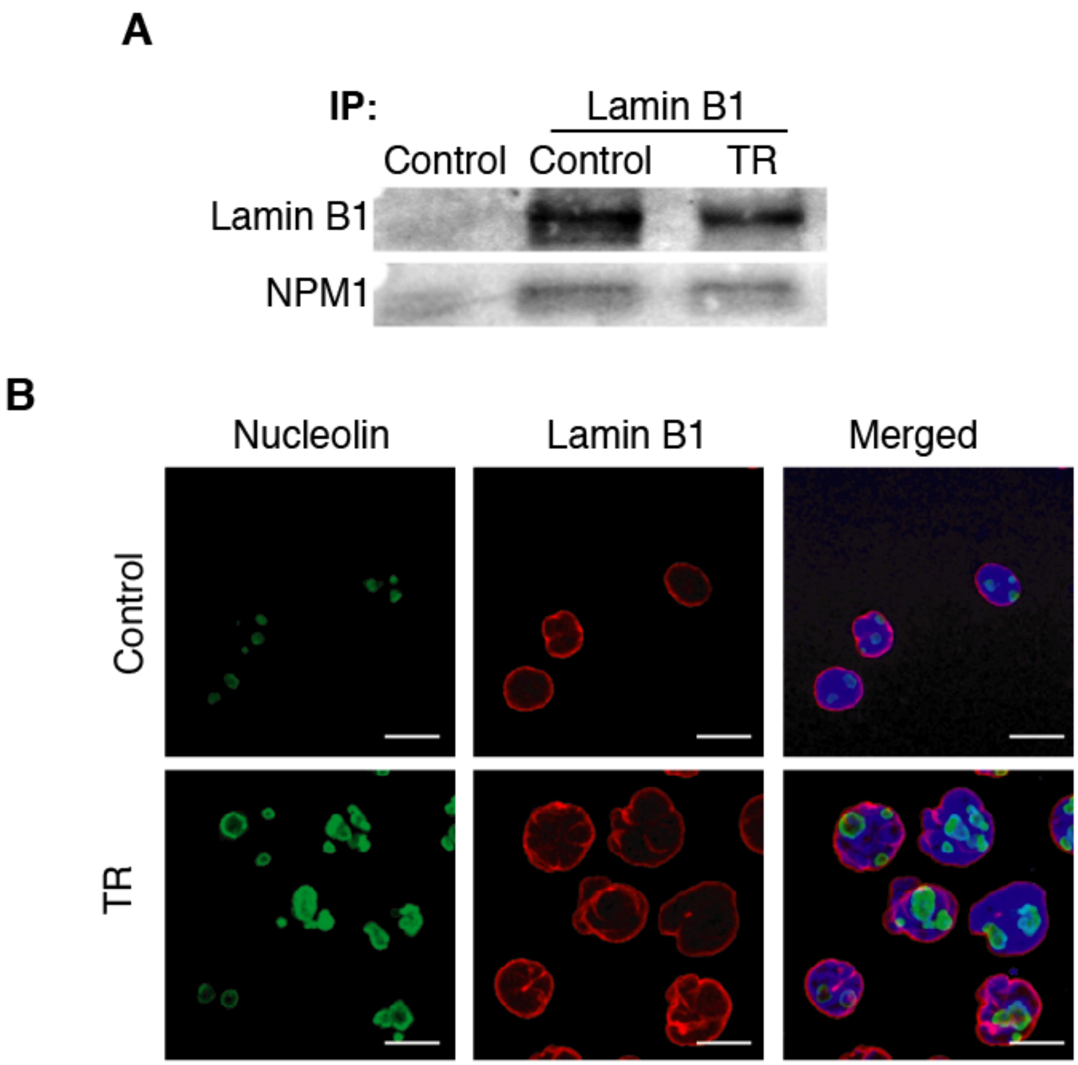


**Figure S2**

**Fig. S2. (A)** Control and TR cells were lysed, sonicated and immunoprecipitated with anti-lamin B1 antibody. The interaction with NPM1 was resolved by immunoblot. **(C)** Control or TR cells were seeded on poly-L-lysine-coated glasses and stained with Hoechst (blue), nucleolin (green) and anti-lamin B1 antibody (red) for their analysis by confocal microscopy. Bar 10 µm.

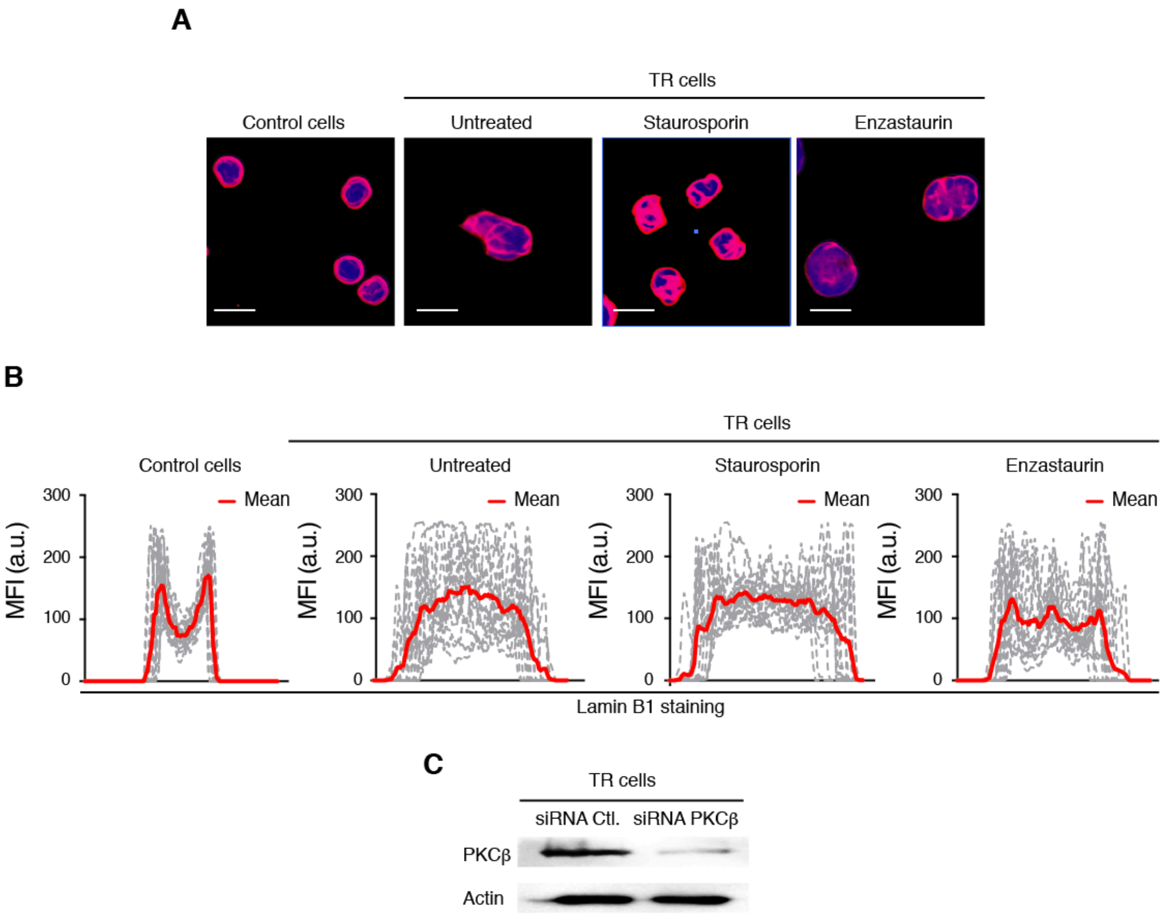


**Figure S3**

**Fig. S3. (A)** Control and TR cells treated or not with 50 nM staurosporin (PKCα inhibitor) or 2 nM enzastaurin (PKCβ inhibitor) for 30 min at 37ºC. Then, cells were seeded on poly-L-lysine-coated coverslips, fixed and stained with Hoechst and anti-lamin B1 antibody. Bar 10 µm. **(B)** Line plots show the signal profile of lamin B1 from 15 representative nuclei. Red line indicates the mean intensity of the profiles analyzed. **(C)** Representative immunoblots for PKCβ and actin in the whole cell lysates of TR cells transfected with a specific pool of siRNAs against PKCβ.

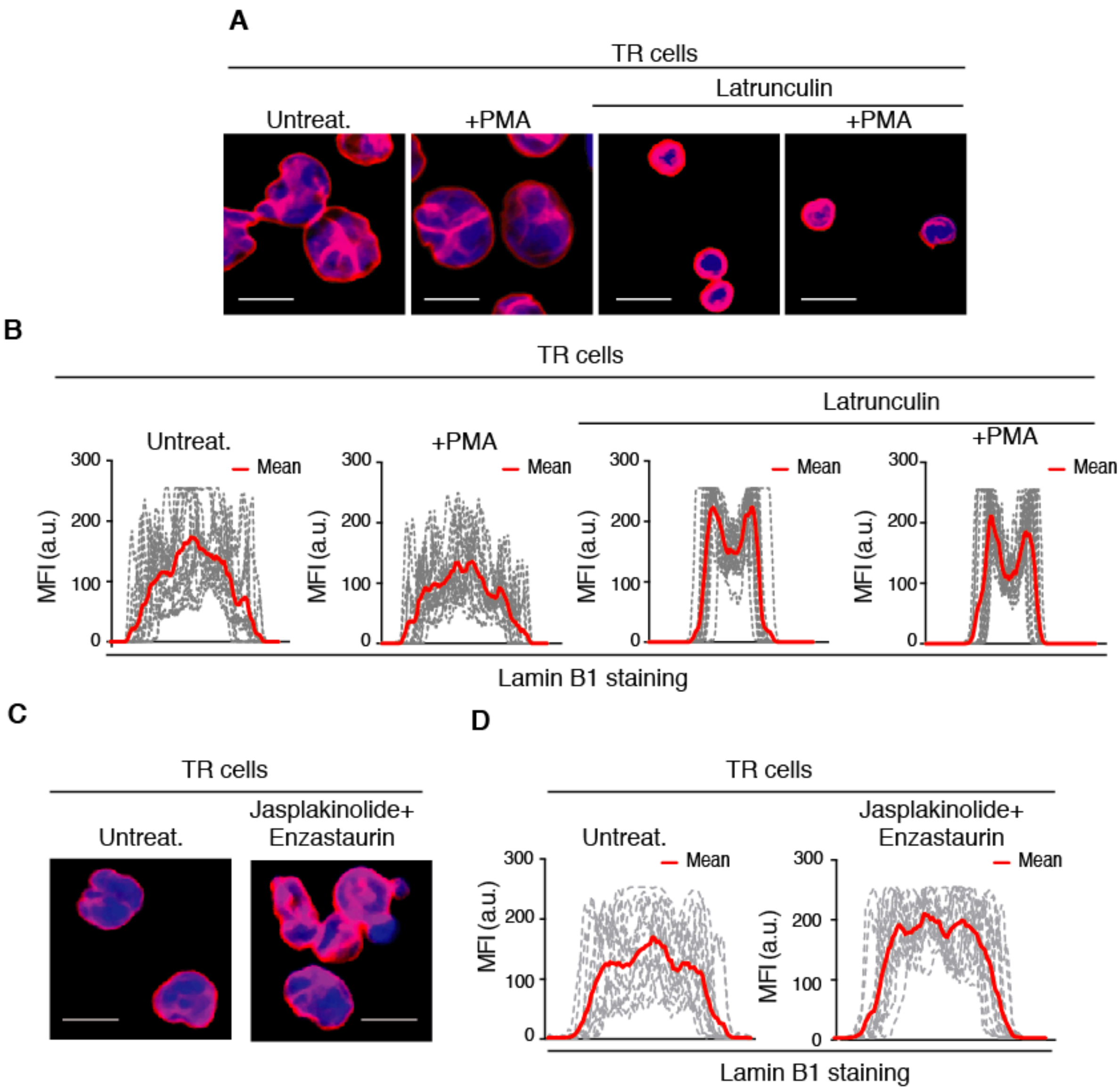


**Figure S4**

**Fig. S4. (A)** TR cells treated or not with latrunculin B (2 µg/mL) for 1 h and were stimulated with PMA (50 ng/mL) for 15 min at 37ºC before seeding on poly-L-lysine coated coverslips. Cells were fixed, permeabilized and stained for their analysis by confocal microscopy. Bar 10 µm. **(B)** Line plots show the signal profile of lamin B1 from 15 representative nuclei. Red line indicates the mean intensity of the profiles analyzed. **(C)** TR cells were preincubated or not with japlakinolide (1 µg/mL for 1 h) and then treated with enzastaurin 2 nM for 30 min before seeding on poly-L-lysine coated coverslips. Cells were fixed, permeabilized and stained for their analysis by confocal microscopy. Bar 10 µm. **(D)** Line plots show the signal profile of lamin B1 from 15 representative nuclei. Red line indicates the mean intensity of the profiles analyzed.

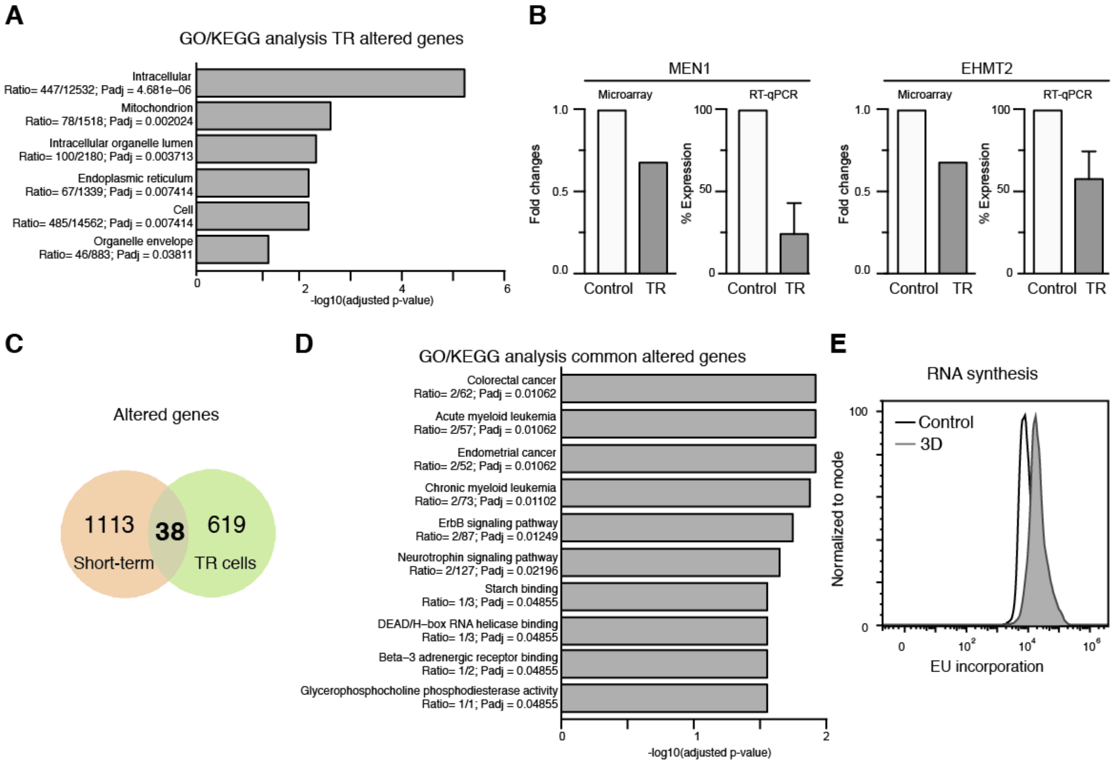


**Figure S5**

**Fig. S5. (A)** Graph shows the top GO/KEGG enrichment results from the transcriptional analysis of control and TR cells by microarray. **(B)** Graph shows the validation of microarray by qPCR. The expression of *MEN1* and *EHMT2* was detected in control and TR cells by qPCR (4 replicates) and the microarray data (2 replicates). Error bars indicate standard deviations. **(C)** Venn diagram representation of common transcriptional changes between short- and long-term 3D confinement conditions. **(D)** GO/KEGG enrichment analysis of common transcriptional changes from (E). **(E)** Control and TR cells were cultured in the presence of EU, fixed, stained and analyzed by flow cytometry.

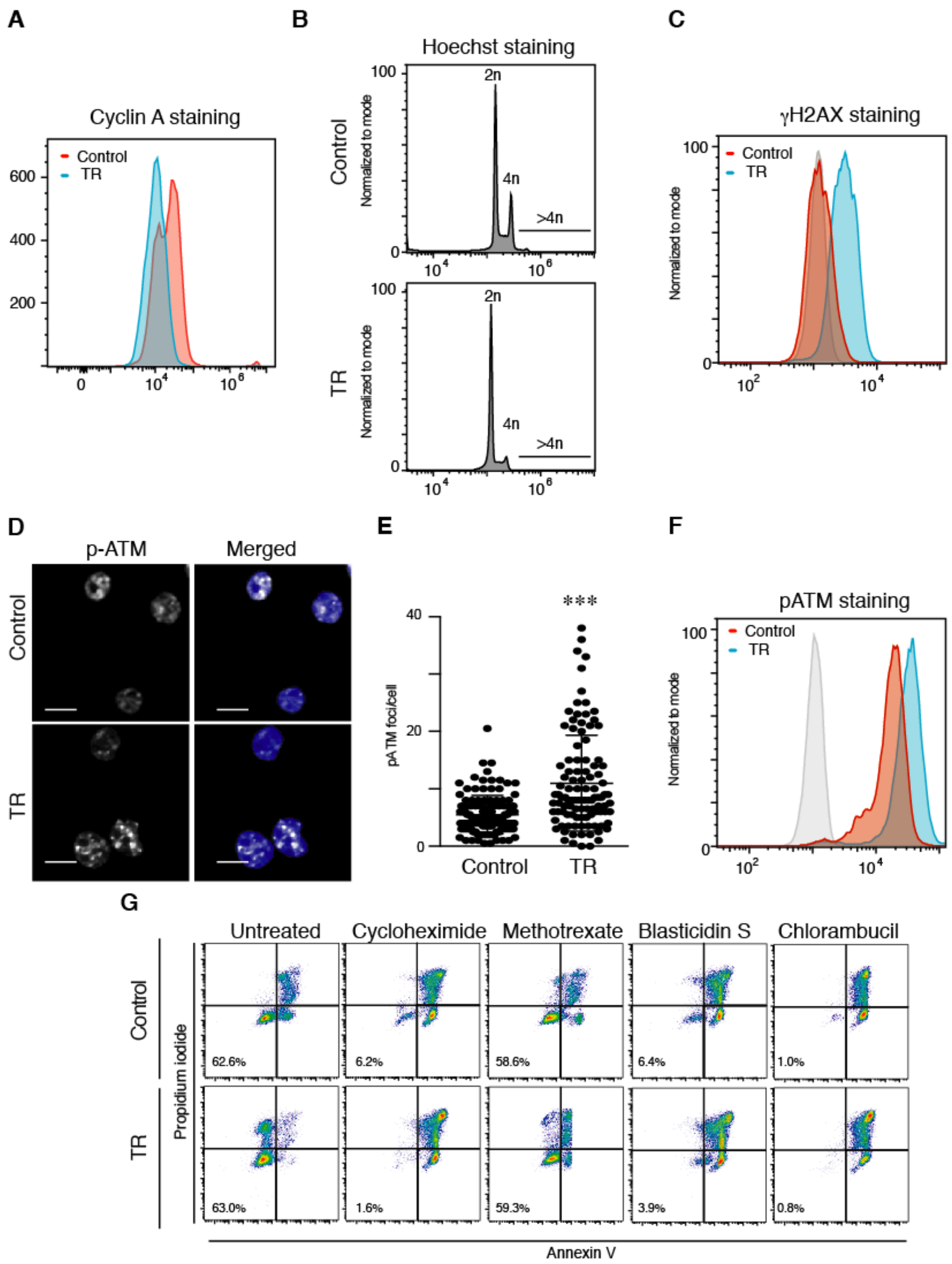


**Figure S6**

**Fig. S6. (A)** Control (red) and TR (blue) cells were fixed, permeabilized and stained with anti-cyclin A antibody. Then, cells were analyzed by flow cytometry. **(B)** Control and TR cells were stained with Hoechst 33342 and ploidy was determined by flow cytometry. **(C)** Control (red) and TR (blue) cells were fixed, permeabilized and stained with anti-γH2AX antibody. Then, cells were analyzed by flow cytometry. **(D)** Control and TR cells were collected, seeded onto poly-L-lysine coated slides, fixed and stained for p-ATM (red) and Hoechst (blue). Bar 10 µm. **(E)** Graph shows the number of foci per nucleus of control or TR cells. n=180-185 (2 independent replicates). **(F)** Control (red) and TR (blue) cells were fixed, permeabilized and stained with anti-p-ATM antibody. Then, cells were analyzed by flow cytometry. **(G)** Control and TR cells were treated with different drugs for 24 h. Then, cells were collected and stained with annexin V-FITC and propidium iodide for their flow cytometry analysis. *** $P < 0.001$.

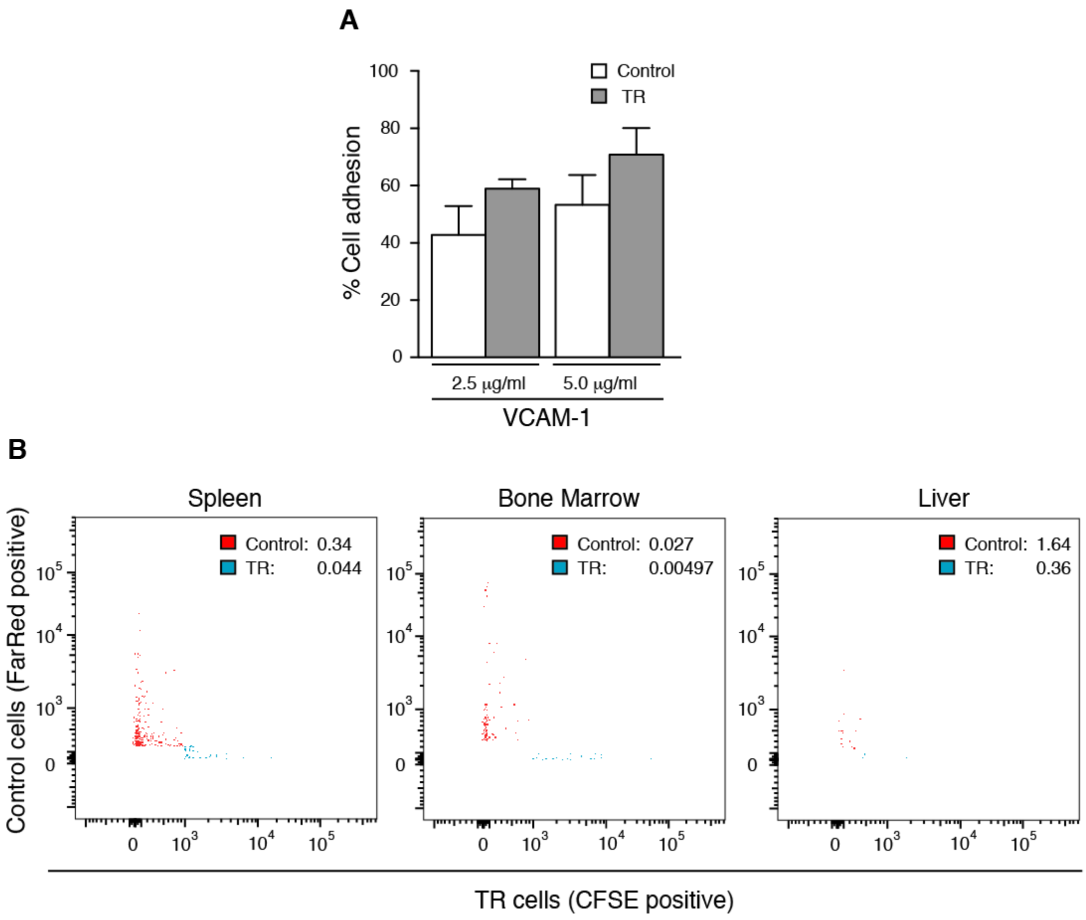


**Figure S7**

**Fig. S7. (A)** Control and TR cells were labeled with 1 µM BCECF-AM (2',7'-Bis-(2-Carboxyethyl)-5-(and-6)-Carboxyfluorescein, Acetoxymethyl Ester) and seeded onto VCAM-1 coated wells at the indicated concentrations. After 30 min, wells were washed with RPMI and attached cells were lysed in PBS-01%SDS and fluorescence was quantified. **(B)** Control and TR cells were labeled with 2 µM Cell Tracker Far Red or 5 µM CFSE respectively, mixed 1:1, and injected into the tail vein of NSG (NOD scid gamma mouse) mice. Graphs show the percentage of labeled cells according to the total number of events in spleen, bone marrow and liver.

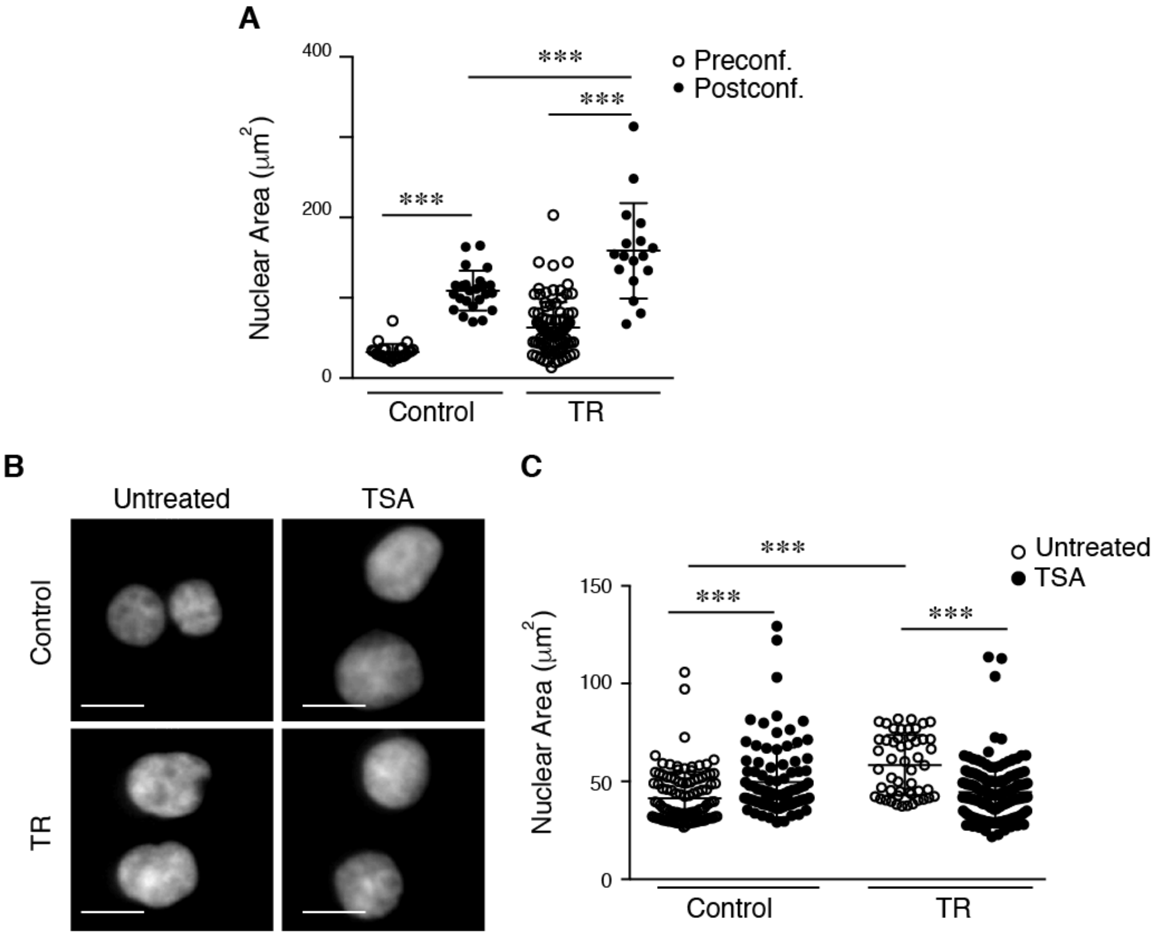


**Figure S8**

**Fig. S8. (A)** Control and TR cells were seeded on poly-L-lysine coated slides, stained with Hoechst, and confined mechanically up to 3 μm. Graph shows the quantification of nuclear area pre- and post-confinement. Mean n=59-121 cells ± SD. **(B)** Isolated nuclei from control and TR cells were seeded on poly-L-lysine-coated coverslips and treated with TSA (300 nM) for 18 h. Then, cells were fixed, and stained with Hoechst. **(C)** Graph shows the nuclear area before (white dots) and after (black dots) TSA treatment in both conditions. Bar 10 μm. *** P < 0.001.

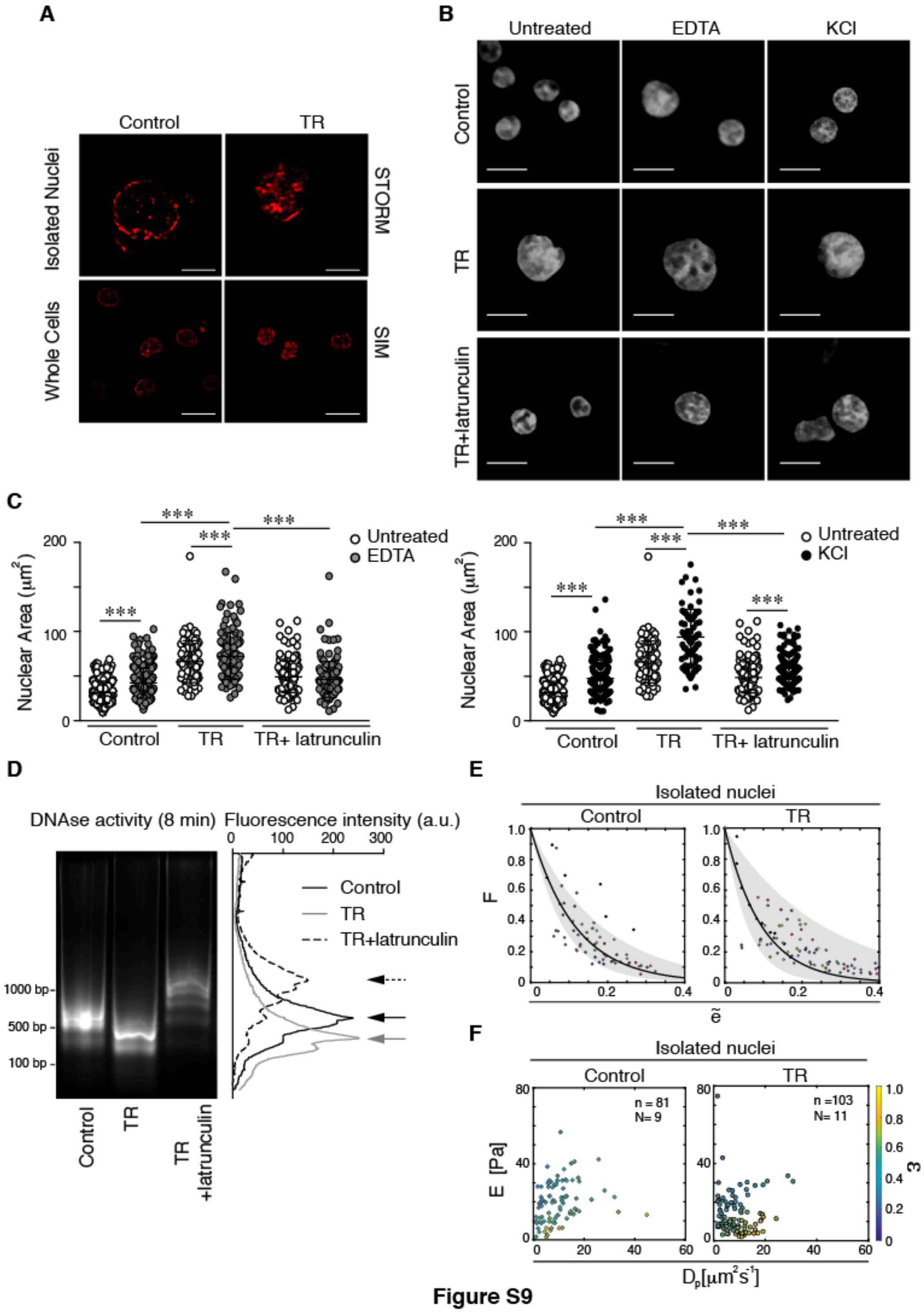


**Figure S9**

**Fig. S9. (A)** Isolated nuclei from control and TR cells were sedimented on poly-L-lysine coated slides and DNA was stained with SiR-DNA (Spirochrome) for 1 h. Then, high resolution images were taken by STORM (Stochastic Optical Reconstruction Microscopy) microscopy Bar 10 μm. Whole control and TR cells were stained with SiR-DNA (Spirochrome) for 1 h and viewed by SIM (Structured Illumination Microscopy) microscopy. Bar 10 μm. **(B)** Isolated nuclei from control and TR cells were seeded on poly-L-lysine-coated coverslips and treated with 5 mM EDTA and KCl for 10 min to induce nuclear swelling. Then, nuclei were fixed and stained with Hoechst for confocal analysis. **(C)** Graphs show the nuclear area quantification upon EDTA (left panel) or KCl (right panel) swelling conditions. Mean n = 1326 nuclei ± SD (2 independent replicates). **(D)** Control and TR cells were incubated with latrunculin B (2 µg/mL) for 1 h before collecting them and digesting their DNA with DNAse for 8 min. Left panel shows the DNA digestion profile from control cells (black line), TR cells (grey line) or TR cells treated with latrunculin B (dashed line). **(E)** Variation of the compaction parameter $\Phi$ with the indentation depth in terms of scaled strain ($\varepsilon$) for isolated nuclei of control and TR cells. Data from each indentation assay are displayed in different colours. The exponential average fitting is shown as a solid black curve and the grey shadowed region represents the standard deviation of the fitting. **(F)** Representation of the correlation between stiffness (E) and force relaxation characteristic time ($\tau_p$) expressed as poroelastic diffusion coefficient ($D_p \sim \delta R/\tau_p$) for isolated nuclei of control and TR cells. Indentation depth in terms of $\varepsilon$ is represented as a color scale. Data points (n) and number of assays (N) per condition are specified. *** $P < 0.001$
